# Searching across working memory and perception

**DOI:** 10.64898/2026.09.15.751716

**Authors:** Dengxinyi Wei, Daniela Gresch, Anna C. Nobre

## Abstract

Searching for objects of interest is an essential and pervasive activity in adaptive cognition. In everyday situations, we often search for relevant content in our surroundings and in mind at the same time, yet most experimental tasks consider search behavior in perception or in working memory separately. To address the gap, we designed tasks to investigate perceptual and working-memory search proceeding in tandem. Participants viewed successive arrays of two real-world objects, separated by a delay, and then were prompted to search for a specific object, while the second array remained visible. The probed target was equally likely to be in the previously encoded array, available only in working memory, or in the visible array. Behavioral performance across four experiments with different response demands and delays yielded a consistent pattern of findings. Accuracy was better for detecting perceptual targets than working-memory targets. However, search speed was equivalent for targets in both domains in all cases, suggesting parallel search functions co-existing across the two domains. The final experiment combined continuous color reproduction with eye tracking and neural recordings. Neural measures of posterior alpha lateralization, tracking the neural selection of working-memory and perceptual targets, suggested parallel search functions developing with similar time courses. Intriguingly, oculomotor gaze shifts showed a divergent pattern, with earlier gaze shifts associated with perceptual targets and later shifts with working-memory targets. Overall, the present studies establish a means for investigating search across working-memory and perceptual domains, suggest the co-existence of parallel search mechanisms, and invite new lines of enquiry.

## Introduction

Attention guides our search for relevant items in the external, sensory environment and in internal, mental representations (Chun et al., 2011; Nobre & Gresch, 2025; van Ede & Nobre, 2023). Of these two domains, search during external perception has been studied most extensively, thus providing the primary basis for our understanding of attentional selection during search. Since the earliest studies (Egeth et al., 1972; Treisman & Gelade, 1980), research has identified many factors guiding and constraining performance for search in the external environment (Bar, 2004; Duncan & Humphreys, 1989; Eckstein, 2011; Wolfe, 2021). For the internal domain, recent studies have begun to reveal parallels between factors that guide and constrain memory search and visual search, suggesting that internal and external selection might rely on some common principles (Kong & Fougnie, 2019, 2022).

Although most research has examined internal and external search separately, searching in real- world situations often combines both internal and external domains in tandem. For example, when baking a cake, we search through and focus on memorized ingredients while simultaneously searching for them in the external environment of the kitchen. In these types of searches, internal and external representations typically serve different roles. In external visual search, working-memory templates often specify the target, while the sensory stream contains the contents for searching. In contrast, in internal visual search, sensory stimuli often guide the search through working-memory contents.

However, very few studies have explored how search unfolds across both domains together, when the target may occur either in working memory or in the sensory environment. As a notable example, Oliva and colleagues (2004) employed a panoramic-search task to compare performance when finding objects that were currently visible or hidden from view. The study highlighted the pragmatic choice of searching through memory or sensory contents (cf. Kumle et al., 2025). Participants strongly preferred searching visible items despite having reliable, well-learned memory representations, but could search memory effectively when required. A more recent study combined working-memory and perceptual search within trials (Liu & van Ede, 2025). After participants encoded two differently colored objects in working memory, a colored cue presented alongside a four-item greyscale array indicated which working-memory item was the search target in the sensory array. Intriguingly, using gaze biases as a proxy for internal and external target selection (cf. van Ede, 2026), they found similar time courses for initiating search through the working-memory and the perceptual array.

The present study builds on existing research efforts to investigate the everyday situation of searching in tandem through contents held in mind and those that are perceptually available. Our new task design places working-memory and perceptual contents on equal footing for search, with the target equally likely to be in either domain. Across four experiments, participants viewed two successive arrays containing two real-world objects. The first was encoded into working memory, whereas the second remained visible when a probe defined the search target, which could be present in either array with equal probability. Participants searched across these internal and external sets to identify the target (Experiment 1), report its location (Experiments 2 and 3), or reproduce its color (Experiment 4). Experiment 4 additionally used eye tracking and electroencephalography (EEG) to compare the temporal dynamics of selecting targets from internal or external arrays.

Across all experiments, responses to internal and external targets were equally fast, although external targets yielded greater accuracy and precision. Posterior alpha lateralization (Experiment 4), a neural proxy for attentional selection (Benedek et al., 2014; Rihs et al., 2007; Wallis et al., 2015; Worden et al., 2000), showed similar timing but was stronger for selection of external targets. Interestingly, the gaze biases accompanying target selection followed a different temporal pattern. Fixational biases consistently emerged earlier for external than for internal targets. Overall, the findings reveal highly efficient coordination when searching for contents across both internal and external domains, with matched behavioral and neural timing but distinct oculomotor dynamics.

## Experiment 1

### Methods

#### Participants

Experiment 1 and all following experiments adhered to the ethical principles of the Declaration of Helsinki. All procedures were approved by the Institutional Review Board of Yale University.

The experiment was conducted online, hosted on Pavlovia (https://pavlovia.org/). To determine the appropriate sample size, we conducted a power analysis assuming a paired *t*-test, with Cohen’s *d* = 0.5, α = 0.05, and power = 0.85. This analysis indicated a required sample size of approximately 38 participants to detect a moderate effect size. The number was calculated with the R package ‘pwr’ (https://github.com/heliosdrm/pwr), which provides functions for power analysis following Cohen (2009). We recruited a larger sample to account for participant exclusion due to increased variability and reduced task engagement that are common in online experiments. We collected data from 51 participants. Eleven participants were excluded based on our initial criteria, which required sufficient task engagement and applied an a-priori behavioral trial-removal procedure (see ‘Data analysis’). All remaining 40 participants (age range: 19 to 40; mean age: 32.03; gender: 15 female, 25 male; handedness: 35 right-handed, 5 left-handed) reported having normal or corrected-to-normal vision. Electronic consent was obtained before individuals started the study. Participants were paid $12 per hour for their participation.

#### Stimuli and apparatus

The task was programmed using PsychoPy3 Experiment Builder (v2024.1.5; Peirce et al., 2019). The different phases of the experiment used a set of 16 unique real-object stimuli selected from a published object databank (Brady et al., 2008). Each of the 16 objects could be presented in 16 different colors. The 16 colors were systematically sampled to have an equal distance (22.5° apart) on a 360° color wheel (Bae et al., 2015). The colors were grouped into four sets so that each color within a group had a 90° difference from its neighboring colors to maximize distinguishability between objects within each trial (Figure 1A). For example, color 1 at 0° was grouped with color 5 (90°), color 9 (180°), and color 13 (270°). Similarly, color 2 at 22.5° was grouped with color 6 (112.5°), color 10 (202.5°), and color 14 (292.5°). Hence, there were four valid color combinations: (1) colors 1, 5, 9, and 13; (2) colors 2, 6, 10, and 14; (3) colors 3, 7, 11, and 15; (4) colors 4, 8, 12, and 16 (Figure 1B).

**Figure 1.**
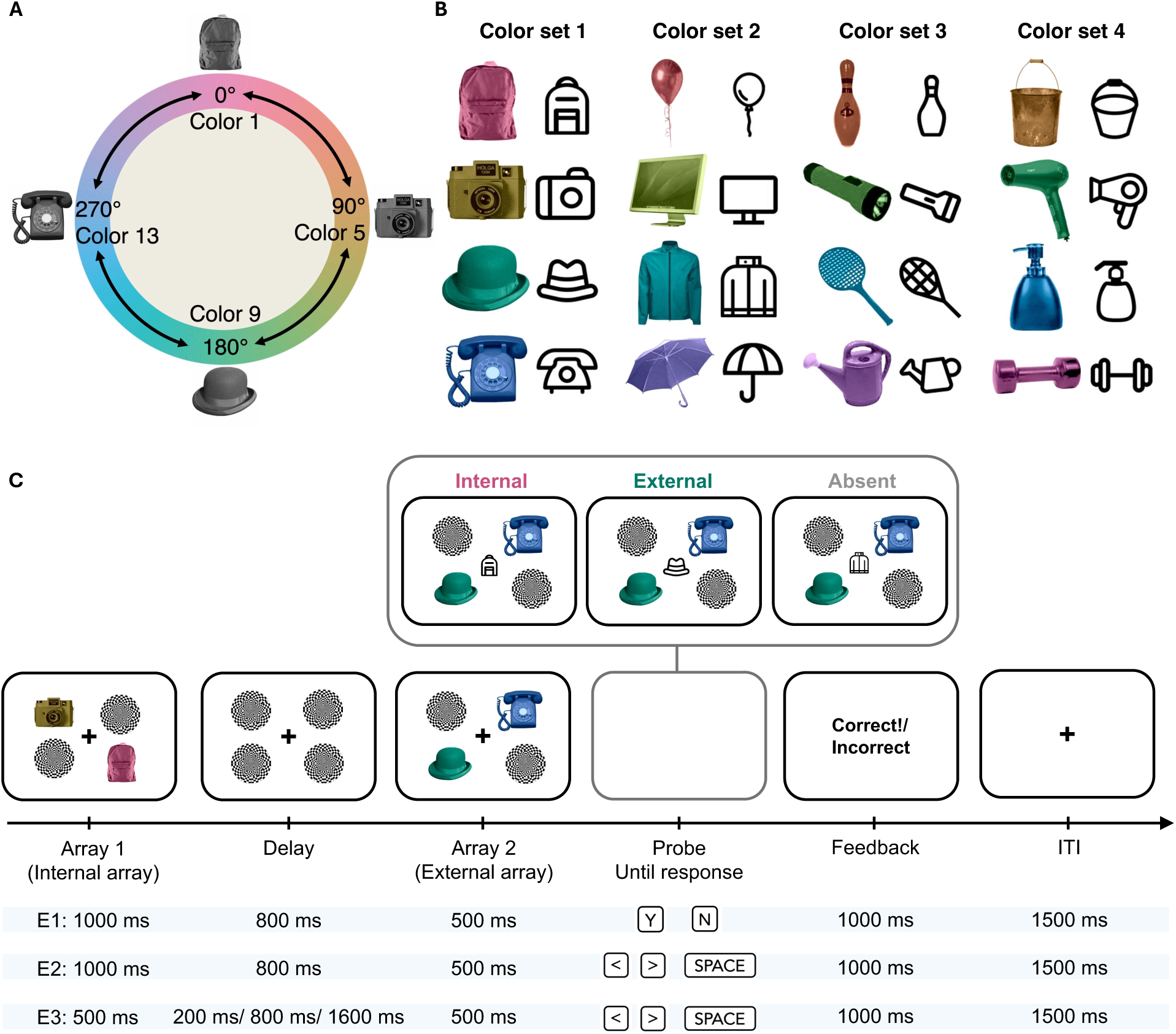
Real-world objects and their corresponding icons used across all experiments, and the schematic of the search task in behavioral Experiments 1, 2, and 3. (A) In each trial, four colors 90° apart from one another were selected. The color combination 1 (out of 4 possible combinations) is illustrated here. (B) There were 16 real-world objects and 16 associated icons. For each object, 16 colors were systematically sampled at equal distances on a 360° color wheel (22.5° apart). Sixteen colors (all illustrated here) were superimposed on gray-scale objects to create 256 colored real-world stimuli. One color is shown for each object as an example. Each column corresponds to one valid color combination (“set”). (C) Blue-highlighted lines indicate the temporal structure and response type in each experiment. First, two objects appeared in the internal array and were subsequently obscured by masks during the delay. Two additional objects then appeared in the previously unoccupied locations. After 500 ms, the fixation cross was replaced by an icon indicating the target object participants were required to search for in either the internal or external array. The external array remained on the screen until participants responded. In Experiment 1, participants were instructed to press “Y” if the target object appeared in the trial, either in the internal or external array, and to press “N” if the target object was absent. In Experiment 2 and Experiment 3, participants reported the location of the identified target by pressing the left or right arrow key. If the target was absent, they pressed the spacebar. The timings of the initial internal array and the subsequent delay differed across experiments. The initial array duration was longer in Experiments 1 and 2 (1000 ms) than in Experiment 3 (500 ms). The delay was fixed in Experiments 1 and 2 (800 ms) and varied across three durations in Experiment 3 (200, 800, 1600 ms).

In addition, greyscale objects were created for presentation during the training phase of the experiment. Complementing the real-world objects, 16 corresponding icons were manually selected from a repository for SVG icons (https://www.svgrepo.com/) and modified using Adobe Illustrator (Figure 1B). Apart from objects and icons, an image of a round black-and-white checkerboard served as a mask. The black fixation cross (RGB value: [0, 0, 0]) and all stimuli were presented against a white background (RGB value: [255, 255, 255]).

Stimulus sizes and locations were scaled with Psychopy “height” units, such that all dimensions were defined relative to the height of the display and therefore remained consistent across different screen resolutions. This yielded a vertical coordinate range of −0.5 to 0.5, with horizontal extent determined by screen aspect ratio (e.g., ±0.8 for a 16:10 display). In the training task, stimulus sizes were defined as follows: objects (0.20; i.e., 20% of screen height), icons (0.10), and fixation cross (0.02). Stimuli were presented above or below the central fixation at positions (0, 0.15) and (0, -0.15). For the search task, sizes were objects (0.15), icons (0.10), masks (0.16), and the fixation cross (0.02). Stimuli were presented either at fixation (0, 0) or at the four quadrant positions: top-left (-0.168, 0.168), top-right (0.168, 0.168), bottom- left (-0.168, -0.168), and bottom-right (0.168, -0.168). Assuming a viewing distance of 50 cm on a 14-inch 16:10 monitor, object and mask stimuli subtended approximately 3.5° of visual angle and were presented at an eccentricity of ∼5 DVA, while icons subtended ∼2 DVA and were always presented at fixation.

#### Task and procedure

The experiment consisted of a training phase followed by the main task. The training phase included a learning task and a memory-test task. In the training phase, participants learned and were tested on the associations between objects and their corresponding icons. The main experiment was an object-search task. The aim was to compare object search when the target could be either in an array encoded into working memory or in a perceptually available array. The search was initiated by the presentation of an icon corresponding to the real-world target object. At the beginning of the experiment, participants were instructed and reminded to maintain central fixation throughout the task, however, eye position was not monitored.

##### Training phase

The training consisted of a learning task and a memory-test task. In the learning task (Supplementary Figure S1A), object-icon pairings were presented. In each trial, one of the 16 objects appeared in greyscale paired with its corresponding icon. The object was presented above the fixation cross, whereas the icon was presented below it. Each object-icon pair appeared on the screen for 1000 ms, followed by a 500-ms inter-trial interval (ITI). Participants were instructed to view the pairs. All object-icon pairings were presented in random order once in each of four blocks, for a total of 64 trials.

In the memory-test task, pairs of corresponding or non-corresponding greyscale objects and icons were presented. The same locations as in the preceding learning phase were used. Participants had to decide whether the object-icon pair was corresponding or non-corresponding (Supplementary Figure S1B). Participants pressed the “Y” key to indicate a match and “I” key for a mismatch. Visual feedback (500 ms) was immediately provided after each response, displaying “Correct!” for a correct identification and “Incorrect” for an incorrect one. After feedback, a 500-ms ITI preceded the next trial. Sixteen matching and 16 non-matching object-icon pairs were intermixed and presented in random order in each of two blocks, for a total of 64 trials.

##### Main experiment

The object search task was of primary interest in our investigation. Participants performed a combined working-memory and perceptual search task in which they searched for a target object that was equally likely to be present in the working-memory array or in the perceptual array of that trial (Figure 1C, E1).

At the start of each trial, an array containing two colored objects and two masks appeared for 1000 ms surrounding the fixation cross. The display was divided into four quadrants, with objects centered within two of the quadrants and masks occupying the remaining two. Objects were placed in one of six possible spatial configurations: (1) top-left and bottom-right, (2) top-left and bottom-left, (3) top-left and bottom-right, (4) bottom-left and top-right, (5) bottom-left and bottom-right, and (6) top-right and top-left. Spatial configurations were pseudo-randomly selected on each trial. This initial display served as the working-memory (i.e., internal) array. During the subsequent 800-ms delay, the two objects were replaced by masks presented at all four quadrant locations. Access to the object identities for search within the initial array at the end of the trial thus relied on internal representations maintained in working memory. After the delay, a second array was presented for 500 ms, in which two objects appeared in the two quadrants that had been previously unoccupied in the initial internal array (i.e., external array). For both the internal and external displays, the identities of the objects were pseudo-randomly drawn from the object pool to avoid repetition throughout the experiment. Object colors were drawn randomly from one of the pre-defined color sets, with the distribution of color sets counterbalanced across the experiment to minimize color-related bias.

Next, the central fixation changed into an icon, while the objects in the second array remained perceptually visible and were therefore externally available for search. The icon indicated the target object for search on that trial. Participants indicated whether the target was present in the trial by pressing “Y” for present or “N” for absent. When present, the target could be either in the internal, working-memory array or in the external, perceptual array. The target was equally likely to originate from the internal or external array and to appear in any of the four quadrants. After each response, participants received visual feedback (i.e., “Correct!” or “Incorrect”) presented at fixation for 500 ms. The ITI was 1500 ms.

The experiment contained 432 search trials, divided into eight blocks of 54 trials. In total, there were 144 (33%) present internal targets, 144 present external targets, and 144 absent targets. Before starting the main search task, participants completed 25 practice search trials (eight present-internal, eight present-external, and nine absent). The entire experiment, including training, lasted about 60 minutes.

#### Data analysis

Data were analyzed in R Studio (Posit team, 2022). Since the main experiment required strong object-icon associations, datasets with an average accuracy of 80% or lower in the training-phase test task were removed (*n* = 1).

In the search task, trials with reaction times (RTs) longer than 5000 ms or shorter than 150 ms were removed. We additionally removed trials with RTs exceeding ±3 times the interquartile range relative to each participant’s performance across all conditions (present-internal, present-external, or absent). Moreover, we removed the entire block of trials if accuracy in any of the search-task conditions (present- internal, present-external, absent) within that block was below 50%. Datasets for which more than 10% of the search trials were rejected were removed from further analysis (*n* = 10). After these exclusion steps, datasets from 40 participants remained in the main analysis, with an average of 98.15% (*SD* = 0.02) of trials retained.

Performance in the training phase was not central to the research question but served to ensure that participants could reliably identify the search target based on its associated icon. Accordingly, when reporting performance in the memory-test task, we present only descriptive statistics (*M* ± *SD*) for accuracy and RTs from correct trials.

For the main search task, the primary comparison of interest was between the two levels of target domain for target-present trials (internal vs. external). Consequently, the analyses and results focus primarily on comparisons between internal and external targets, while the target-absent condition is reported more succinctly in the Supplementary Materials (see ‘Supplementary Materials’). When reporting performance in the search task, we present descriptive statistics (*M* ± *SD*) for accuracy and RTs from correct trials, along with the results of inferential analyses. We conducted paired *t*-tests comparing the internal and external conditions, reporting Cohen’s *d* as a measure of effect size.

## Results

### Training

Participants successfully learned the correct object-icon correspondences. Overall performance was high for both match (accuracy = 0.98 ± 0.03; RT = 633.16 ± 188.61 ms) and mismatch trials (accuracy = 0.97 ± 0.03; RT = 693.09 ± 195.86 ms; Supplementary Figure S1C, first row).

### Search

Reaction times from 40 participants were submitted to a paired *t*-test to assess differences between the levels of target domain (internal vs. external) in target-present trials. Results showed that the mean RTs for identifying targets in the external array (886.14 ± 258. 21 ms) and internal array (868.52 ± 251.98 ms) did not differ significantly, (*t*(39) = 1.85, *p* = 0.07, *d* = 0.29; Figure 2A, left panel). For accuracy, participants were less accurate when targets were present in internal arrays (0.89 ± 0.08) compared to external arrays (0.96 ± 0.03; *t*(39) = 5.36, *p* < 0.001, *d* = 0.86; Figure 2A, right panel). We also tested whether performance differed between target-present and target-absent trials. Participants responded more slowly when the target was absent from the arrays than in target-present trials, whereas search accuracy did not differ between conditions (Supplementary Table 1A, Supplementary Figure S2).

**Figure 2.**
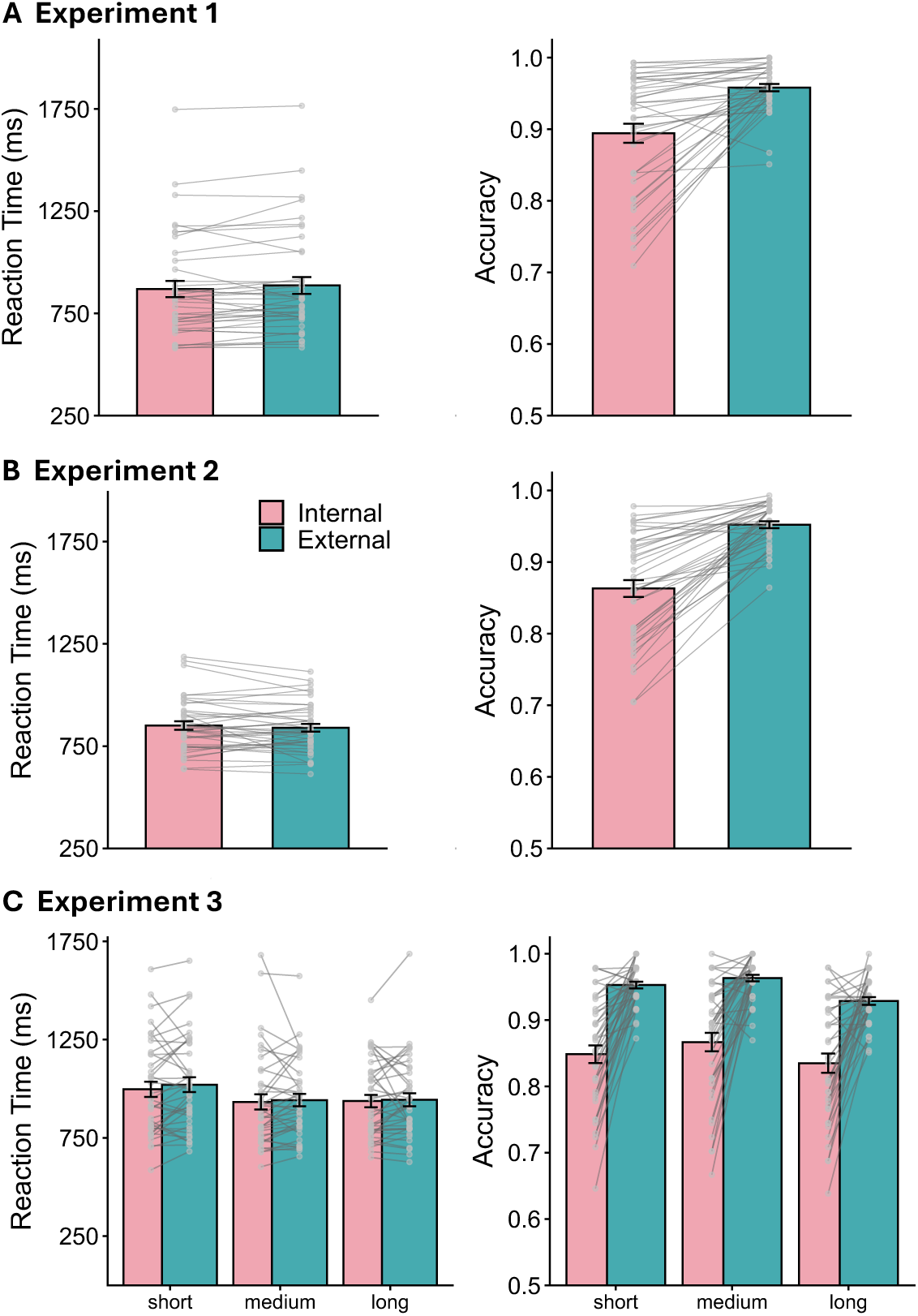
Search-task results for Experiments 1, 2, and 3. The left panel shows RTs, and the right panel shows accuracy. Gray dots indicate the mean RT and accuracy for each participant. Error bars indicate standard errors of the mean (*SEM*). (A) In Experiment 1 (*n* = 40), RTs for identifying targets in the internal and external arrays did not differ significantly. Accuracy was lower for targets in the internal array than for targets in the external array. (B) In Experiment 2 (*n* = 41), RTs did not differ significantly between targets in the internal and external arrays. Accuracy was significantly lower for targets in the internal array than for targets in the external array. (C) In Experiment 3 (n = 39), RTs for identifying targets in the internal and external arrays did not differ significantly, whereas short delays between the two arrays resulted in longer RTs. Across all delay durations, accuracy was lower for targets in the internal array than for targets in the external array. Accuracy was highest when the delay between arrays was of medium duration.

In summary, Experiment 1 yielded two new observations. Objects in working memory were acted upon as quickly as perceptually available objects on the screen. However, performance accuracy for identifying internal objects was diminished compared to identifying external objects.

## Experiment 2

Experiment 2 tested whether the pattern of results observed in Experiment 1 generalized to task conditions that additionally required spatial localization of the search target rather than simple present-absent identification. Using the same stimulus parameters and overall procedures, participants searched for targets either in the internal, working-memory array or the external, sensory array. When the target was present, they reported its corresponding hemifield (left or right). In contrast to Experiment 1, in which spatial location was not required for the response, accurate performance in Experiment 2 relied on both identity and spatial location of the target.

### Methods

#### Participants

Given its similarity to Experiment 1, we planned a sample size of *n* = 38 to detect significant effects (*p* = 0.05, two-tailed) with a power of 0.85. Online participants were recruited via Prolific and Yale University’s SONA platform (https://yale.sona-systems.com/). Similar to Experiment 1, we recruited a larger sample to account for participant exclusion. We collected data from 67 participants. Twenty-six participants were excluded based on the initial exclusion criteria (see ‘Data analysis’). All remaining 41 participants (age range: 18 to 33; mean age: 21.02; gender: 18 female, 23 male; handedness: 30 right-handed, 9 left-handed, 2 ambidextrous) had normal or corrected-to-normal vision. Electronic consent was obtained before individuals participated in the study. Participation via SONA provided undergraduate students with course credit (1 credit per hour), whereas Prolific participants were paid $12 per hour for their participation.

#### Stimuli and apparatus

The stimuli and apparatus were identical to those in Experiment 1.

#### Task and procedure

Experiment 2 closely adhered to the procedures of Experiment 1. The training phase was identical to that in Experiment 1. The search phase differed from Experiment 1 only in relation to the reporting requirements of the search task. When an icon appeared to prompt the search target, participants reported the target’s location (left or right hemifield) by pressing the corresponding arrow key if the target was present in either the internal or external array. If the target was absent, participants pressed the space bar (Figure 1C, E2).

The experiment included 432 visual search trials, divided into eight blocks of 54 trials. In total, there were 288 present targets, equally divided into four conditions (internal-left, internal-right, external-left, and external-right; 72 per condition). The target was absent in 144 trials.

#### Data analysis

Data were preprocessed according to the pipeline described for Experiment 1. In Experiment 2, no participants were removed for low accuracy in the memory-test task. In the search task, RT- and accuracy-based exclusions removed 26 participants. After these exclusion steps, datasets from 41 participants remained in the main behavioral analysis, with an average of 98% of trials retained (*SD* = 0.01).

As in Experiment 1, the training phase was completed to train the correspondence between objects and icons. For the main search task, we applied the identical data analysis procedure as for Experiment 1.

### Results

#### Training

Overall performance was high for both match (accuracy = 0.97 ± 0.04; RT = 706.30 ± 211.21 ms) and mismatch (accuracy = 0.98 ± 0.04; RT = 746.92 ± 210.48 ms) trials, suggesting participants successfully learned the correct target-icon correspondences (Supplementary Figure S1C, second row).

#### Search

Reaction times from 41 participants were submitted to a paired *t*-test to assess differences between the levels of target domain (internal vs. external) in target-present trials. Results showed the mean RTs for identifying targets in the external array (840.02 ± 118.88 ms) and internal arrays did not differ significantly (850.78 ± 31.36 ms; *t*(40) = 0.99, *p* = 0.33, *d* = 0.16; Figure 2B, left panel). For accuracy, participants were less accurate when targets were presented in internal arrays (0.86 ± 0.07) compared to external arrays (0.95 ± 0.03; *t*(40) = 9.11, *p* < 0.001, *d* = 1.42; Figure 2B, right panel). Moreover, we tested whether performance differed depending on whether the target was present versus absent in the arrays. Same as in Experiment 1, when the target was absent, participants were consistently slower, but search accuracy did not differ between the two conditions (Supplementary Table 1B; Supplementary Figure S2).

The pattern of findings replicated that observed in Experiment 1, suggesting that the results generalize to task conditions in which target reporting relies on both identity and spatial location.

## Experiment 3

Experiment 3 was designed to investigate the time course of search in working memory versus perception. We systematically varied the interval between the internal and external arrays, thereby manipulating the time available to encode and maintain the internal representations before the external array appeared. At the shortest interval (200 ms), the internal array would likely still be represented in iconic memory when the external array appeared. Although access to iconic representations is typically high (Sperling, 1960), the presentation of the external array may interfere with the encoding of the internal contents into working memory. At the intermediate (800 ms) and longest intervals (1600 ms), the internal array is likely maintained in working memory, with different opportunities for decay. Internal search performance could improve with longer intervals if additional time supports encoding and maintenance of the internal items before the external array appears. Alternatively, internal search performance could decline as representations transition from iconic to working memory and become increasingly susceptible to decay over time.

Furthermore, although search accuracy for targets from the internal domain was lower than for targets from the external domain, mean accuracy remained high in both Experiments 1 and 2, and RTs did not differ significantly between target domains. Therefore, in Experiment 3, we not only manipulated the interval following the internal array but also shortened its presentation duration to 500 ms to match that of the external array. This modification served two purposes. First, it equated the duration of “pure” visual input for internal and external objects before the probed target appeared, thereby rendering the temporal structure of both arrays more comparable. Second, it increased the demands of internal search by reducing the time available to encode the working-memory objects, allowing us to test whether limiting encoding time would impair performance.

### Method

#### Participants

We aimed for the same sample size as in Experiments 1 and 2 (*n* = 38) to detect significant effects between internal and external conditions (*p* = 0.05, two-tailed) with a power of 0.85. Participants were recruited online using Prolific and Yale SONA. We collected data from 68 participants. Twenty-nine participants were excluded following the same preprocessing procedure as in Experiments 1 and 2 (see ‘Data analysis’). All remaining 39 participants (age range: 19 to 40; mean age: 32.46; gender: 21 female, 18 male; handedness: 34 right-handed, 3 left-handed, 2 ambidextrous) had normal or corrected-to-normal vision. Electronic consent was obtained before individuals could participate in the study. Participation via SONA provided undergraduate students with course credit (1 credit per hour), whereas Prolific participants were paid $12 per hour for their participation.

#### Stimuli and apparatus

The stimuli and apparatus were identical to those in Experiments 1 and 2.

#### Task and procedure

Experiment 3 closely adhered to the procedures of Experiment 2. The training phase was identical to that in Experiments 1 and 2.

For the search phase, the only difference concerned the temporal structure of the trial. The internal array was presented for 500 ms and was followed by a short (200 ms), medium (800 ms), or long (1600 ms) delay (i.e., delay duration). The medium delay matched that used in Experiments 1 and 2. After the delay, the external array appeared, and the fixation cross was replaced by an icon. Participants reported the location of the probed target (left or right hemifield), regardless of whether it was present in the internal or external array. When the target was present, they pressed the corresponding left or right arrow key. If the target was absent, they pressed the space bar (Figure 1C, E3).

**Figure 3.**
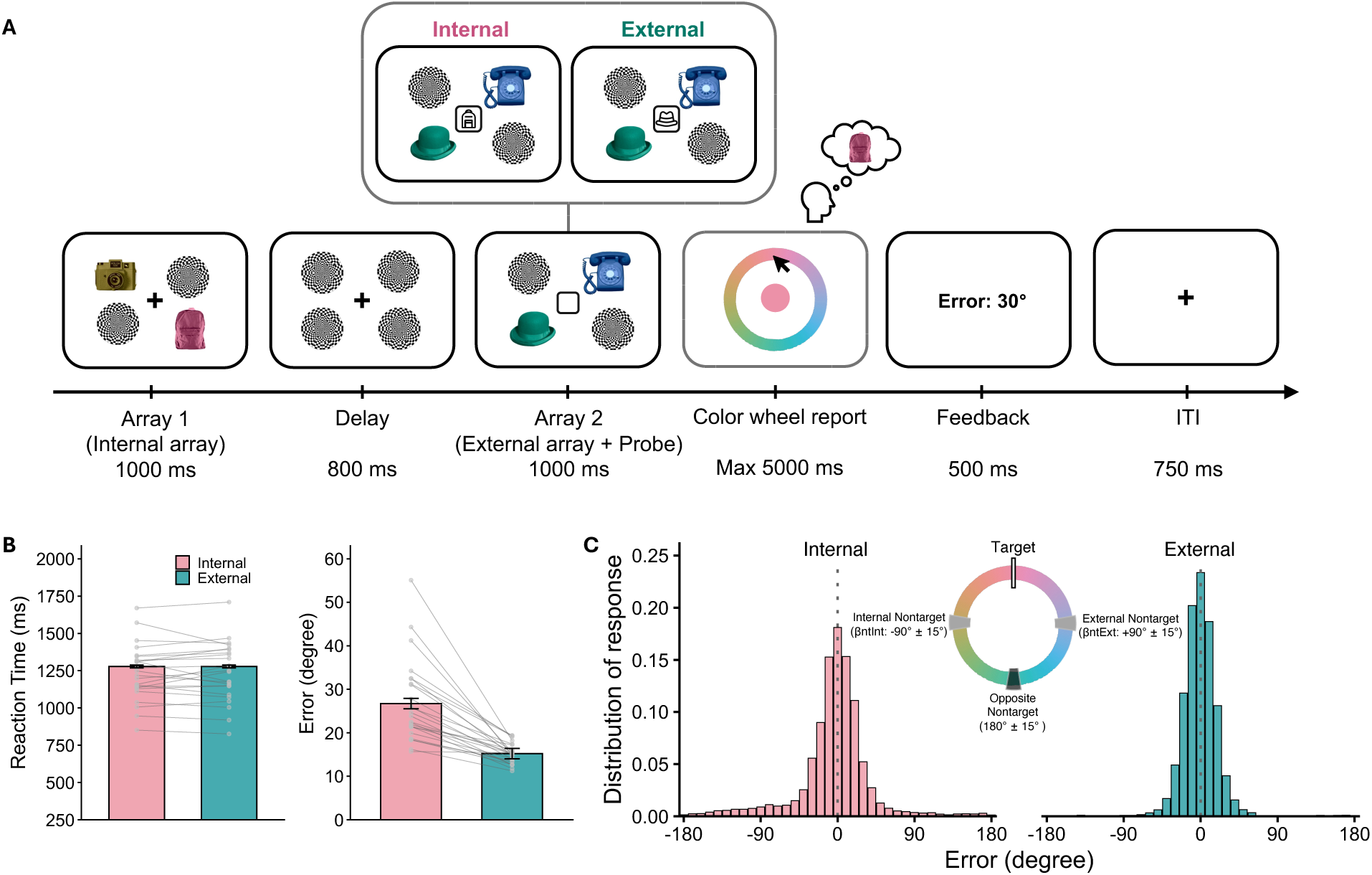
Schematic of the search task in Experiment 4 and behavioral results. (A) Unlike in Experiments 1-3, the external array was presented simultaneously with the probe, and participants responded using a continuous color wheel report. The icon in Array 2 was surrounded by a black square to provide a uniform visual boundary across different icons. Numerical feedback was presented to indicate the accuracy of participants’ color reports relative to the target color. The illustration here shows the participant reporting the color of an internal target. (B) In Experiment 4 (*n* = 25), RTs did not significantly differ between targets presented in the internal and external arrays. Error reports were significantly higher for targets presented in the internal array than for those presented in the external array. (C) Group-level response-error distributions for internal-target reports (pink) and external-target reports (teal). Errors are plotted relative to the target color, aligned to 0°. The internal nontarget was aligned to -90° (±15°) and the external nontarget to +90° (±15°), as illustrated by the color wheel. Internal-target reports were broader and included more off-target responses, whereas external-target reports were more tightly centered around the target.

Experiment 3 included 432 visual search trials, divided into eight blocks of 54 trials. In target-present conditions (288), trials were equally distributed across internal and external targets, left or right hemifields, and the three delay-duration conditions (24 trials each). Trials in the target-absent conditions (144) were equally divided across the short, medium, and long interval conditions (48 trials per condition).

#### Data analysis

Data were preprocessed according to the pipeline described for Experiments 1 and 2. In Experiment 3, one participant was removed based on low accuracy on the memory-test task. In the search task, RT- and accuracy-based exclusions removed 28 participants. After these exclusion steps, datasets from 39 participants remained in the main behavioral analysis, with an average of 97.8% (*SD* = 0.01) of trials retained.

Similar to Experiments 1 and 2, the training phase ensured participants could reliably identify the target object based on its associated icon. Thus, we report only descriptive statistics (*M* ± *SD*) of accuracy and RTs for correct trials.

For the search task, the primary conditions of interest were target domain (internal vs. external) and delay duration (short vs. medium vs. long) for target-present trials. We therefore conducted repeated-measures analyses of variance (ANOVAs) on RT and accuracy data. We report partial eta squared (η^2^_p_) as a measure of effect size. When the assumption of sphericity was violated, Greenhouse-Geisser-corrected *p*- values (*p*[GG]) are reported. For post-hoc *t*-tests, we report Bonferroni-corrected *p*-values and Cohen’s *d*.

### Results

#### Training

Overall performance was high for both match (accuracy = 0.98 ± 0.02; RT = 664.01 ± 151.71 ms) and mismatch (accuracy = 0.97 ± 0.03; RT = 717.40 ± 157.66 ms) trials, suggesting participants successfully learned the correct target-icon correspondences (Supplementary Figure S1C; third row).

#### Search

Reaction times from 39 participants were submitted to a two-way repeated-measures ANOVA with target domain (internal vs. external) and delay duration (short vs. medium vs. long) as factors. Results showed a significant main effect of delay duration (*F*(2, 76) = 20.78, *p*[GG] < 0.001, η^2^_p_ = 0.35), but no significant main effect of target domain (external: 968.59 ± 95.73 ms; internal: 955.88 ± 99.13 ms; *F*(1, 38) = 0.67, *p* = 0.42, η^2^_p_ = 0.02) or significant interaction (*F*(2, 76) = 0.38, *p*[GG] = 0.67, η^2^_p_ = 0.01). Post-hoc comparisons for the effect of delay duration indicated that participants were slower in trials with a short delay compared with a medium delay (*t*(38) = 5.62, *p* < 0.001, *d* = 0.90) or a long delay (*t*(38) = 5.38, *p* < 0.001, *d* = 0.86). However, RTs did not differ between medium and long delays (*t*(38) = 0.24, *p* = 1.00, *d* = 0.04; Figure 2C, left panel). For accuracy, the results showed a significant main effect of target domain (*F*(1, 38) = 75.1, *p* < 0.001, η^2^_p_ = 0.66), a significant main effect of delay (*F*(2, 76) = 9.72, *p* < 0.001, η^2^_p_ = 0.20), but no interaction between target domain and delay (*F*(2, 76) = 0.03, *p*[GG] = 0.71, η^2^_p_ = 0.01). Post-hoc pairwise comparisons with Bonferroni correction revealed that participants were less accurate when targets were present in the internal array (0.85 ± 0.03) compared to the external (0.94 ± 0.01; *t*(38) = 8.67, *p* < 0.001, *d* = 1.39). For the effect of delay duration, post-hoc comparisons indicated higher accuracy in medium-delay trials than for long-delay trials (*t*(38) = 3.92, *p* = 0.001, *d* = 0.63). However, accuracy did not differ between short and medium delays (*t*(38) = 2.34, *p* = 0.07, *d* = 0.38), or between short and long delays (*t*(38) = 2.4, *p* = 0.06, *d* = 0.38; Figure 2C, right panel).

Moreover, we tested whether performance differed depending on whether the target was present versus absent. As in Experiments 1 and 2, when the target was absent, participants were consistently slower, but the search accuracy did not differ between the two conditions (Supplementary Table 2; Supplementary Figure S2).

In Experiment 3, we replicated the pattern of results observed in Experiments 1 and 2. Search times for identifying and localizing targets in internal and external arrays were equivalent. This pattern persisted despite the shortened presentation time of the internal array and remained stable across all delay durations between the two arrays. Accuracy was also consistently lower when localizing target objects in internal arrays compared to external arrays. Thus, across the three experiments, the patterns of RTs and accuracy were highly consistent (Supplementary Figure S3), indicating strong generalizability of the observed effects.

## Experiment 4

Experiment 4 was conducted in person and combined behavioral measures, eye tracking, and EEG recordings. The behavioral experiment was designed to provide a finer-grained assessment of search performance across working memory and perception. Participants used a precision response to reproduce the color of the target object on a continuous color wheel. The introduction of the continuous color wheel report required adaptations to the task design. To separate search-related activity from response-related activity, the external array and the probe were presented for 1000 ms before participants could initiate their response and then disappeared at the response stage. This provided a period during which neural responses and fixational gaze behavior could be examined without contamination from the response itself. The timing of the perceptual search display was also adjusted. Unlike in Experiments 1-3, the external array and probe were presented simultaneously. This ensured that search began before a stable representation of the external items had been formed, allowing us to examine neural and gaze responses at the onset of perceptual search.

### Methods

#### Participants

The target sample size was set at *n* = 25. This decision was informed by the robust within-participant effects observed across Experiments 1-3 and by previous in-person studies from our laboratory using comparable working-memory, continuous-report, oculomotor, and neural measures (Boettcher et al., 2021; Gresch, Boettcher, van Ede, et al., 2024). Across Experiments 1-3, the internal-versus-external accuracy contrast yielded absolute paired-samples effect sizes of Cohen’s *d* = 0.85, *d* = 1.42, and *d* =1.39, respectively. For Experiment 4, we adopted a conservative planning effect equal to 75% of the smallest previously observed effect, corresponding to *d* = 0.64. An a priori power analysis conducted using the R package ‘pwr’ indicated that a two-sided paired-samples test with alpha = 0.05 and power of 0.85 required 25 participants.

Participants were recruited through Yale University’s BrainWorks SONA (https://yale-wtibrainworks.sona-systems.com/) and physical poster advertisements in New Haven, CT. We collected data from 28 participants. Three participants were excluded prior to analysis due to excessive EEG noise observed during recording. All remaining 25 participants (age range: 19 to 32; mean age: 24; gender: 18 female, 6 male, 1 non-binary; handedness: 24 right-handed, 1 left-handed) had normal or corrected-to-normal vision. Electronic consent was obtained before individuals could participate in the study. Participants were paid $30 per hour for their participation.

#### Stimuli and apparatus

Participants were seated 106.5 cm away from a 69-cm GIGABYTE M27Q monitor (1,920 x 1,080-pixel resolution; 60-Hz refresh rate). An EyeLink 1000 Plus eye tracker (SR Research), positioned approximately 15 cm in front of the monitor, recorded binocular gaze continuously at a sampling rate of 1000 Hz.

EEG data were acquired using a Brain Products actiCHamp Plus amplifier and recorded with the Lab Streaming Layer (LSL) Connector and LSL Recorder software. Signals were acquired from 64 scalp electrodes arranged according to the international 10-10 system, together with two auxiliary channels recording horizontal and vertical electrooculography (HEOG and VEOG). During the EEG setup, impedances were reduced to below 5 kΩ where possible and maintained below 10 kΩ for all electrodes. Horizontal and vertical eye movements were monitored using bipolar EOG recordings, with one electrode pair placed lateral to the outer canthus of each eye (HEOG) and another pair positioned above and below the left eye (VEOG). For visualization, the data were referenced online to the Fz electrode. During acquisition, the EEG and EOG signals were digitized at a sampling rate of 1000 Hz and stored for offline analysis.

The experiment was programmed in Python using the PsychoPy package (v2024.1.5; Peirce et al., 2019). We used the same set of 16 unique real-object stimuli and icons also used in Experiments 1-3. On each trial, four colors were systematically sampled from a 360° color wheel. A base color was first selected randomly from the color wheel, and the remaining three colors were sampled at offsets of +90°, +180°, and +270° relative to the base color, with the hue values wrapping around the color wheel when exceeding 360°. To increase color variability across trials and reduce potential categorical color effects, each sampled color was independently jittered by ±15°. Consequently, the angular separation between adjacent colors ranged from 60° to 120°. For example, if the base color was located at 1°, the remaining three colors would be centered at 91°, 181°, and 271°, with each color independently jittered by ±15°.

In the training task, stimulus sizes and locations were identical in size to those in Experiments 1-3. For the search task, stimulus sizes and positions were specified in units of visual angle using PsychoPy’s ‘deg2pix’ conversion based on the monitor geometry. Colored object stimuli and masks were 2.5° of visual angle (∼150 × 150 pixels). Probe icons were 0.83° of visual angle (∼50 × 50 pixels). The fixation cross was 0.5° of visual angle (∼30 pixels), and the enlarged fixation cross was 1° of visual angle (∼60 pixels). The color wheel had an outer diameter of 8.3° of visual angle (∼500 pixels) and an inner diameter of 5.0° of visual angle (∼300 pixels). The feedback text was 0.33° of visual angle (∼20 pixels). Stimuli were presented either at fixation (0°, 0°) or at one of four peripheral locations centered ±4.5° of visual angle from fixation along the horizontal and vertical axes (top-left: -4.5°, 4.5°; top-right: 4.5°, 4.5°; bottom-left: -4.5°, -4.5°; bottom-right: 4.5°, -4.5°), corresponding to ∼ ±270 pixels from the center of the display.

#### Task and procedure

The experiment consisted of a training phase followed by the main task. The training phase was identical to that in Experiments 1-3. Eye-tracking and EEG data were not acquired during this phase because the purpose was simply to familiarize participants with the object-icon associations. Overall performance was high for both match (accuracy = 0.96 ± 0.04; RT = 719.79 ± 224.02 ms) and mismatch (accuracy = 0.98 ± 0.03; RT = 766.87 ± 249.00 ms) trials, suggesting participants successfully learned the correct target-icon correspondences.

In the main experiment, participants performed a combined working-memory and perception search task in which they searched for a target object equally likely to be present in a working-memory array or a perceptual array, before reporting its color (Figure 3A).

At the start of each trial, a bold fixation cross was presented for 250 ms, signaling the beginning of the trial, followed by a standard fixation cross for 500 ms. An array containing two colored objects and two masks was then presented for 1000 ms around the fixation cross. The display was divided into four quadrants, with objects appearing in two quadrants and masks occupying the remaining two. Objects were arranged in one of two spatial configurations: (1) top-left and bottom-right or (2) top-right and bottom-left. These configurations were counterbalanced across trials in the experiment. This initial display served as the working-memory (i.e., internal) array.

Following an 800-ms delay, during which masks replaced the objects at all four quadrant locations, a second array was presented for 1000 ms (i.e., external array). Two colored objects appeared in the two previously unoccupied quadrants, with two masks in the other locations. Simultaneously with this external array, an icon was presented at the center, indicating the target object for search. The icon was surrounded by a black square to provide a uniform visual boundary across different icon shapes.

For both the internal and external arrays, object identities were randomly sampled from the object pool, whereas the target object (indicated by the icon) was counterbalanced across the experiment. Object colors were assigned systematically from the color wheel. For each target color, the color approximately 180° away was always assigned to a nontarget object in the opposite domain from the target. Of the two neighboring colors, approximately ±90° from the target, one was assigned to a nontarget object in the same domain as the target, and the other was assigned to a nontarget object in the opposite domain.

Next, a color wheel was presented and remained visible until participants responded or until the 5000-ms response window elapsed. The color wheel was randomly rotated every trial to prevent anticipation of the spatial layout of the colors. Participants reported the color of the probed target object by selecting a location on the color wheel with the mouse. The mouse cursor always started at the center of the screen. The target was equally likely to originate from the internal or external array and to appear in any of the four quadrants. After each response, participants received visual feedback indicating the accuracy of their report (e.g., ‘Error: 30°’), calculated as the angular distance between the reported color and the target color. The possible error ranged from 0° to 180°, with smaller values indicating more accurate reports. The feedback was presented at fixation for 500 ms, followed by a fixed 750-ms ITI.

The experiment contained 512 trials, divided into eight blocks of 64 trials. In total, there were 256 (50%) internal target trials and 256 (50%) external target trials. Before starting the main search task, participants completed 16 practice search trials (eight internal, eight external). The entire experiment, including training, lasted about 90 minutes. Participants could take a break between each block. Prior to each block, the eye tracker was calibrated and validated using a standard five-point procedure. Participants were instructed to maintain central fixation throughout each trial.

#### Data analysis

##### Behavioral analysis

Similar to the data preprocessing pipeline in Experiments 1-3, trials with reaction times (RTs) shorter than 150 ms or longer than 5000 ms were removed. We additionally removed trials with RTs exceeding ±3 times the interquartile range relative to each participant’s performance across all conditions (internal or external). Datasets for which more than 10% of the search trials were rejected were set to be removed from further analysis. After these exclusion steps, datasets from all 25 participants remained in the behavioral analysis, with 98.51% of trials retained overall.

Paired t-tests compared the response times and report errors between target domains (internal vs. external). Descriptive statistics (*M* ± *SD*) for RTs and report errors are provided along with the results of inferential analyses.

For report errors, we further applied a probabilistic mixture model separately to internal- and external-target report conditions to estimate the probabilities of target reports, guesses, and swaps, as well as response precision and report bias, defined as the mean shift in responses (see Narhi-Martinez et al., 2024). The response error was calculated as the signed angular difference between the reported and target colors on the color wheel. Response errors were aligned such that the target color was centered at 0°, yielding errors ranging from -180° to +180°. The alignment was oriented such that the external nontarget was represented at +90° (±15°) and the internal nontarget at -90° (±15°) across all trials. This alignment allowed positive response errors to be interpreted as responses biased toward the external nontarget color and negative response errors as responses biased toward the internal nontarget color.

For each target condition, each participant’s distribution of response errors was fitted with a probabilistic mixture model estimating five parameters: *γ*, the probability of random guessing, modeled as a uniform distribution; κ, the concentration parameter reflecting response precision; μ, the mean shift of the von Mises components; βntInt, the probability of misreporting the internal nontarget, modeled as a von Mises distribution centered at -90°; βntExt, the probability of misreporting the external nontarget, modeled as a von Mises distribution centered at +90°. The probability of reporting the target was estimated as 1 - *γ* - βntExt - βntInt, corresponding to a von Mises distribution centered at 0° plus the estimated mean shift μ (see Formula 1). Reports to the opposite nontarget (180° away from the target) were not modeled as a part of the mixture components.

Model fitting was conducted using a custom maximum-likelihood procedure implemented in R. Parameters were estimated by minimizing the negative log-likelihood using the Broyden-Fletcher-Goldfarb-Shanno (BFGS) optimization algorithm implemented in base R’s ‘optim’ function. This approach is well suited for estimating mixture models in which the mixture weights are constrained to sum to one. Because κ is a unit-free parameter, to better assess the response precision, we derived circular standard deviation (*SD*) in degrees from the κ using the mean resultant length of the von Mises distribution (see Formula 2).

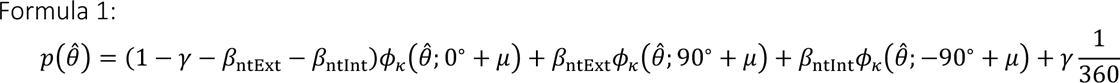

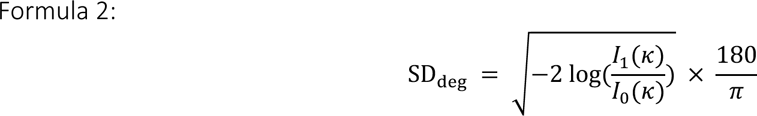

The main comparisons involved target reporting rate, guessing rate *γ*, response precision indexed by *SD*, report bias μ (i.e., mean shift), and swap probabilities βntExt and βntInt. For inferential analyses, planned paired-samples *t*-tests were used to compare internal-target and external-target conditions for target report rate, guessing rate, and SD. We also conducted one-sample *t*-tests on mean shift for internal and external target reports to detect directional biases. Because these comparisons targeted theoretically distinct aspects of performance, including guessing, precision, and bias, *p*-values were reported uncorrected. Swap probabilities were analyzed using a 2 × 2 repeated-measures ANOVA with target domain (internal vs. external) and swap domain (internal nontarget vs. external nontarget) as within-subject factors. We report partial eta squared (η^2^_p_) as a measure of effect size. When the assumption of sphericity was violated, Greenhouse-Geisser-corrected *p*-values (*p*[GG]) are reported. For post-hoc *t*-tests, we report Bonferroni-corrected *p*-values and Cohen’s *d*.

##### Eye-tracking analysis

Eye-tracking preprocessing and analyses followed procedures established in previous work from our laboratory (Draschkow et al., 2022; Gresch, Boettcher, van Ede, et al., 2024; van Ede, Chekroud, & Nobre, 2019; van Ede et al., 2021). Eye-tracking data were converted from EDF to ASC format (SR Support Forum; Eyelink DataViewer: EDF to ASC Conversion tool) and imported into R. Eye blinks were identified and linearly interpolated over a window extending ±100 ms around each blink. Horizontal and vertical gaze positions were obtained by averaging the left and right eye signals. The data were then downsampled to 250 Hz, smoothed using a 60-ms moving average (15 samples), and baseline-corrected by subtracting the mean gaze position during the 200 ms before the second array and target-probe onset from all subsequent time points. Probe-locked epochs were extracted from -200 to 1250 ms relative to target-probe onset.

To focus on fixational gaze biases, trials were excluded if horizontal or vertical gaze deviated by more than ±2.25° of visual angle (∼135 pixels) from central fixation, corresponding to half the distance between fixation and the peripheral stimulus locations (Draschkow et al., 2022; Gresch, Boettcher, van Ede, et al., 2024; van Ede et al., 2021). Participants with more than 20% of trials rejected by this criterion were excluded from further analyses (*n* = 4). The final eye-tracking dataset included 21 participants, with an average trial retention rate of 90% (*SD* = 5.29). While ±2.25° of visual angle seemed like a liberal threshold, the same gaze bias pattern was obtained if we used a stricter threshold (±1° of visual angle (∼60 pixels) from central fixation).

To quantify gaze biases, we first compared the horizontal gaze-position time courses for left- and right-target trials within each target domain. These lateralized gaze traces were then combined into a single measure of gaze bias (i.e., towardness). At each time point, towardness was calculated as the average horizontal gaze position in right-target trials minus that in left-target trials, divided by two. Towardness was used to assess whether gaze was significantly biased toward the target location (i.e., relative to zero) and to compare gaze biases between the internal and external target domains over time. Statistical significance of gaze towardness was evaluated using cluster-based permutation tests implemented in the ‘permuco’ package (Frossard & Renaud, 2021) with 1,024 permutations.

##### EEG preprocessing

EEG data from 25 participants were processed in Python with MNE-Python version 1.10.1 (Gramfort et al., 2013). The data were re-referenced to the common average and assigned the EasyCap M1 electrode montage. Signals were notch-filtered at 60 Hz, band-pass filtered from 0.1 to 50 Hz, and downsampled to 250 Hz. Bad electrodes were identified by visual inspection and interpolated, with an average of 2.84 ± 1.07 electrodes interpolated per participant. Ocular artifacts were removed using independent component analysis (ICA). ICA was fitted to a copy of the continuous EEG data that had been highpass filtered at 1 Hz. Candidate eye-related components were automatically detected with MNE-Python’s ‘find_bads_eog’ function, based on correlations with the recorded HEOG and VEOG channels. These candidate components were then manually evaluated using their scalp topographies, time courses, and EOG correlation scores. Components reflecting blinks or eye movements were removed from the original EEG data. The continuous EEG was epoched from -200 to 1250 ms relative to probe onset, without baseline correction. Although epochs were retained through 1250 ms to cover the partial response period (1000-1250 ms after probe onset), the main analyses focused on the 0-1000-ms post-probe interval. For each epoch, the standard deviation was calculated for each EEG channel and then averaged across channels to yield a single trial-variability estimate. Generalized extreme Studentized deviate (GESD) was applied to these estimates to identify outlier epochs, which were excluded from all subsequent analyses. On average, 13.44 ± 13.69 trials were rejected per participant, corresponding to 2.63 ± 2.67% of trials. Finally, a surface Laplacian transform was applied to improve the spatial resolution of the EEG signals (Boettcher et al., 2021; Gresch et al., 2025; Nasrawi et al., 2023; Perrin et al., 1989; van Ede, Chekroud, Stokes, et al., 2019).

##### Time-frequency analysis

Time-frequency representations of oscillatory power were computed by convolving the EEG data with Morlet wavelets across frequencies from 3 to 40 Hz. The number of cycles increased linearly with frequency (number of cycles = frequency × 0.4), yielding an approximately fixed 400-ms temporal window across frequencies. To quantify visual selection in the internal and external domains, oscillatory power at predefined posterior electrodes (PO7/PO8) was compared between trials in which the target location was contralateral versus ipsilateral to the electrode of interest. Lateralized activity was calculated as a normalized contrast, [(contra - ipsi) / (contra + ipsi)] × 100, and then averaged across the left and right hemispheres. Visual-selection time courses were derived by averaging this lateralization measure across the alpha band (8-12 Hz). Topographical maps were generated by contrasting right- and left-target trials, with normalized differences computed separately at each electrode.

Statistical significance of the contralateral-versus-ipsilateral EEG contrasts was assessed using cluster-based permutation tests (Maris & Oostenveld, 2007). This approach controls for multiple comparisons across time and frequency by evaluating observed clusters against a permutation distribution of the largest cluster statistics obtained after randomly sign-flipping participant-level averages. The cluster-based permutation tests used MNE-Python’s default number of permutations (1,024). A two-tailed cluster-forming threshold of α = 0.05 was used, and clusters with corrected *p* < 0.05 were considered significant. These analyses were applied both to the full time-frequency maps and to time courses averaged within the predefined frequency band of interest.

To compare the timing of alpha lateralization across conditions, we also used a jackknife-based latency analysis (see Gresch et al., 2024). We created 25 leave-one-participant-out grand-average alpha time courses by repeatedly recomputing the group average while excluding one participant at a time. For each leave-one-out waveform, latency was estimated as the time point at which alpha lateralization reached 50% of its peak amplitude. Jackknife standard errors, *t*-statistics, and *p*-values were then computed using the correction procedure described by Miller et al. (1998).

### Results

#### Behavioral results

Reaction times were submitted to a paired *t*-test to assess differences between the target domains (internal vs. external). Results showed that the mean RTs for reporting target colors in the internal array (1277.73 ± 47.80 ms) and external array (1277.94 ± 47.80 ms) did not differ significantly (*t*(24) = 0.02, *p* = 0.98, *d* = 0.003; Figure 3B, left panel). Report errors were also submitted to a paired *t*-test for target domain (internal vs. external). Results showed that the mean error for reporting target colors in the internal array (26.72 ± 5.98°) was significantly greater than the mean error for reporting target colors in the external array (15.21 ± 5.99°; *t*(24) = -6.80, *p* < 0.001, *d* = -1.36; Figure 3B, right panel).

To characterize report performance across working-memory and perceptual targets, we fitted response-error distributions separately for internal- and external-target report trials. Figure 3C shows the distribution of response errors aligned with the target color, with the target at 0°, the internal nontarget at -90° (±15°) and the external nontarget at +90°(±15°).

Figure 4A-C shows the performance for internal and external targets from model-based estimations. Compared with internal-target reports, external-target reports showed a significantly higher probability of target reports (*t*(24) = 5.35, *p* < 0.001, *d* = 1.07), a lower probability of random guessing (*t*(24) = -4.06, *p* < 0.001, *d* = -0.81), and higher precision, reflected by a lower SD parameter (*t*(24) = -9.79, *p* < 0.001, *d* = -1.96). These results indicate that reports were more accurate and precise for targets in the perceptual domain than for targets maintained in working memory, consistent with Experiments 1-3.

**Figure 4.**
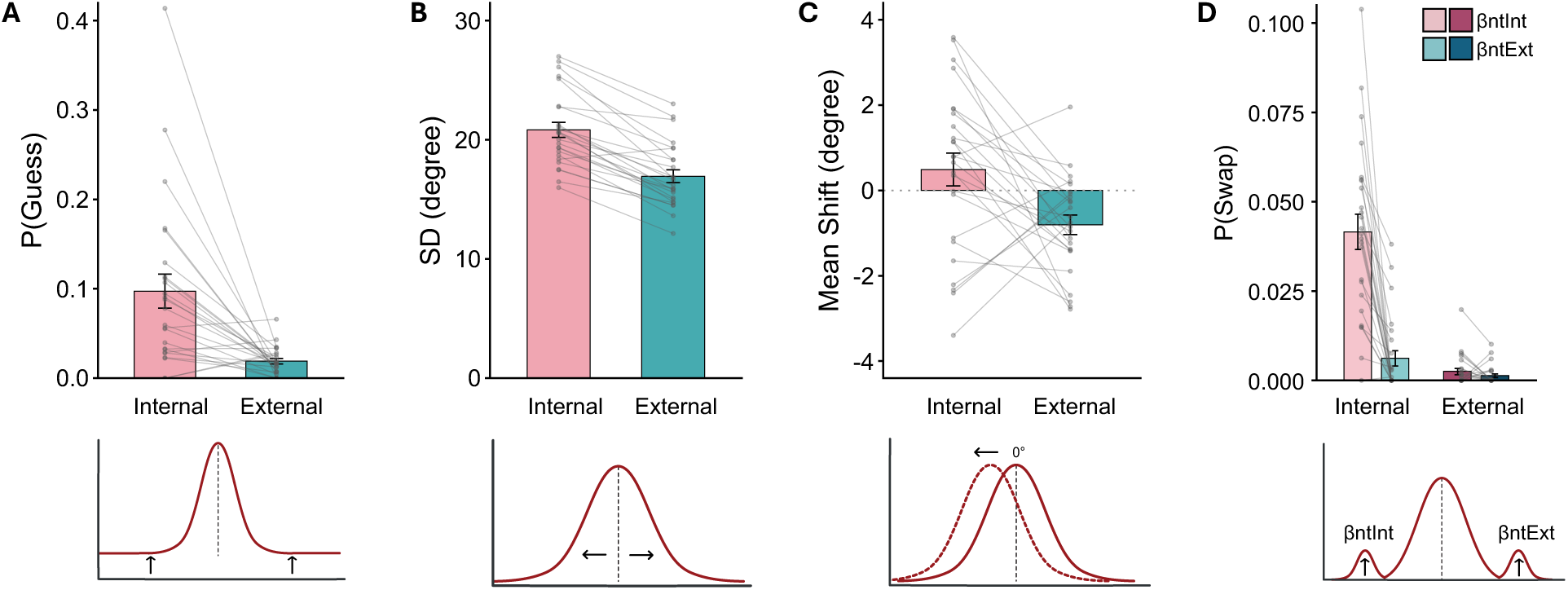
Model-based characterization of response errors for internal- and external-target reports. Parameter estimates from the response-error model, plotted separately for internal- and external-target reports. Bars indicate group means (*M*), black error bars indicate the standard error of the mean (*SEM*), and gray lines show individual participants. Internal-target reports showed a higher probability of guessing (A) and a higher SD (B) than external-target reports, indicating poorer accuracy and precision for memory targets. (C) Internal-target reports showed no significant mean shift, whereas external-target reports showed a significant negative mean shift. (D) Internal-target reports showed elevated swaps with internal nontargets, whereas swap probabilities for external-target reports were low overall.

We next assessed target-report biases by testing whether the mean parameter of the target distribution differed from the true target value of 0° (Figure 4C). One-sample *t*-tests revealed that the mean shift for internal-target reports did not differ significantly from zero (*t*(24) = 1.28, *p* = 0.21, *d* = 0.26). In contrast, external-target reports showed a significant negative mean shift (*t*(24) = -3.53, *p* = 0.002, *d* = - 0.71), indicating that the reports were systematically shifted away from the external nontarget and/or toward the internal nontarget.

Finally, we examined swap errors with the ±90° internal and external nontargets only (excluding 180° opposite nontargets; Figure 4D). A 2 x 2 repeated-measures ANOVA on swap probability, with target domain (internal vs. external) and swap domain (internal nontarget vs. external nontarget) as within-subject factors, revealed significant main effects of target domain (*F*(1, 24) = 65.40, *p* < 0.001, η^2^_p_ = 0.73), and swap domain, *F*(1, 24) = 41.04, *p* < 0.001, η^2^_p_ = 0.63), as well as a significant interaction (*F*(1, 24) = 51.71, *p* < 0.001, η^2^_p_ = 0.68). The results indicate that swap errors were primarily driven by internal-target trials, which were disproportionately vulnerable to swaps with internal nontargets. Bonferroni-corrected follow-up comparisons showed that, for internal-target reports, the swap probability was significantly higher for internal than external nontarget items (*t*(24) = -6.82, *p* < 0.001, *d* = -1.36). By contrast, for external-target reports, the swap probability did not differ between nontarget items from the internal and external domains (*t*(24) = -1.47, *p* = 0.153, *d* = -0.29).

#### Gaze biases

To investigate the oculomotor responses related to search across working memory and perception, we focused our analysis on fixational gaze behavior relative to the probe onset (Figure 5A). To quantify lateral gaze biases, we combined the right- and left-target gaze time courses into a single “towardness” metric for each target domain (Figure 5B). Cluster-based permutation analyses revealed a significant cluster for internal-target trials (∼716-1000 ms, *p* < 0.001; pink horizontal line) and a significant cluster for external- target trials (∼336-776 ms, *p* < 0.001; teal horizontal line), indicating reliable gaze biases toward the target location in both domains.

**Figure 5.**
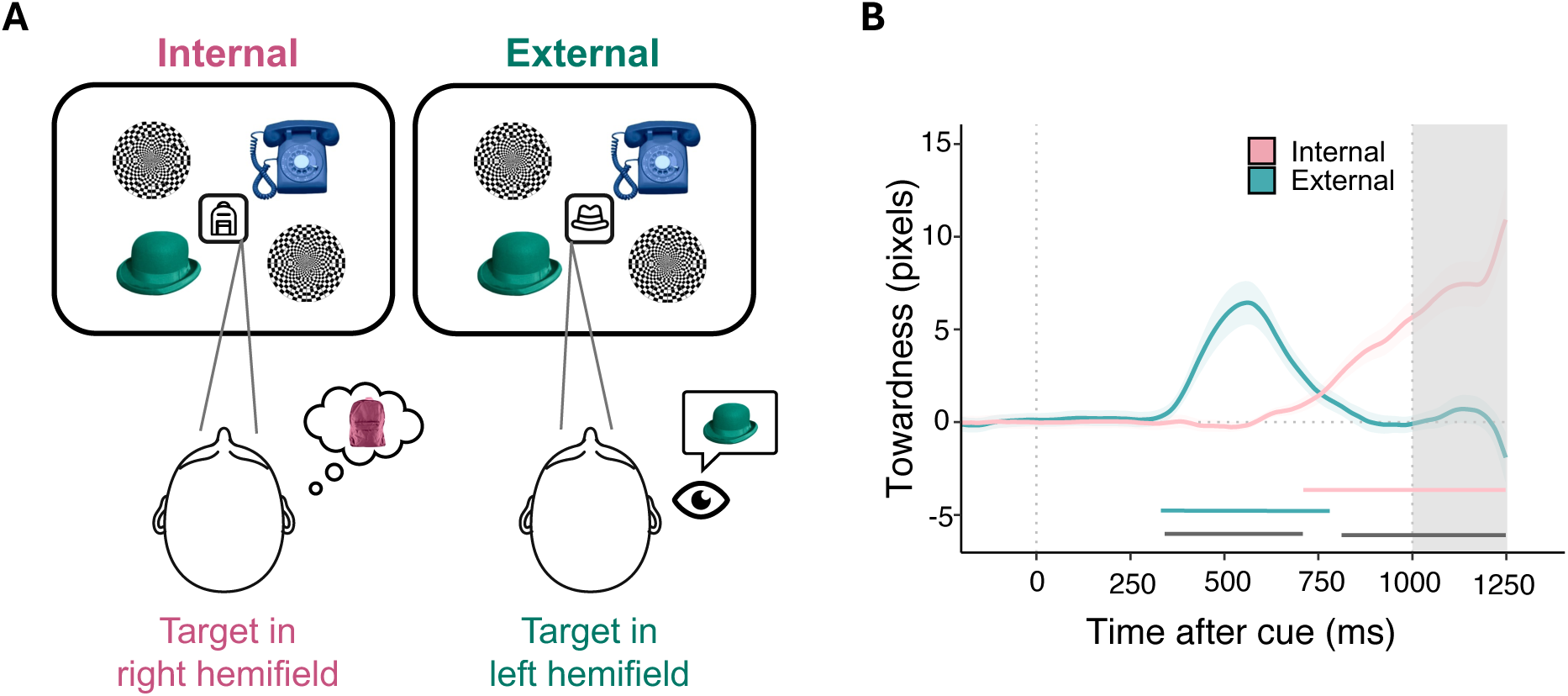
Fixational gaze biases toward target locations in internal and external search. (A) Schematic illustration of gaze biases toward the relevant hemifield in internal- and external-target trials. In internal trials, the probe directed gaze toward the location of an item maintained in working memory; in external trials, the probe directed gaze toward a currently visible item. In the examples shown, the internal target is in the right hemifield, and the external target is in the left hemifield, but these locations varied from trial to trial. (B) Gaze towardness, computed by combining left- and right-target trials into a single measure of gaze bias toward the target location, for internal-target trials (pink), external-target trials (teal), and the difference between internal- and external-target trials (gray). Cluster-based permutation tests revealed significant towardness clusters for both internal and external targets. Gaze biases toward internal targets emerged later than those toward external targets. Shadings indicate *M* ± *SEM*.

The towardness metric also enabled a direct comparison of the time courses of attentional orienting between the internal and external target domains. This comparison revealed two significant clusters (∼348-704 ms, *p* < 0.001; ∼861-1000 ms, *p* < 0.001), indicating that the magnitude of gaze bias differed between the two domains. The time courses suggested that gaze biases toward external targets emerged earlier than those toward internal targets, with little temporal overlap.

#### Time frequency analysis

Posterior alpha-band lateralization is a well-established marker of visual-spatial orienting in both internal and external domains, reflected in differences in 8-12 Hz alpha power over posterior electrodes contralateral versus ipsilateral to the attended location (Benedek et al., 2014; Rihs et al., 2007; Wallis et al., 2015; Worden et al., 2000). We therefore used posterior alpha lateralization as a neural index of spatial selection to ask three related questions: first, whether lateralized alpha activity was present for both working-memory and perceptual selection when participants needed to search across both domains; second, whether the magnitude and temporal patterns of this activity differed between domains, for example whether one signal was more transient or sustained; and third, whether the onset latency of lateralized alpha differed between internal and external selection, indicating whether these selection signals evolved together or followed distinct temporal dynamics.

Consistent with previous work, we observed clear contralateral-versus-ipsilateral attenuation of posterior alpha-band activity following the selection of either internal or external targets. In the alpha-band time-course analysis, selecting internal targets elicited a significant cluster of lateralized alpha attenuation from 536 to 996 ms after probe onset (time-frequency representation: *p* = 0.001; time course: *p* = 0.002). Selecting external targets elicited a more sustained significant cluster from 452 to 1000 ms (time-frequency representation: *p* = 0.001; time course: *p* = 0.001). The corresponding alpha-band time courses showed that selecting internal targets produced a weaker and more transient alpha-lateralization response, whereas selecting external targets produced lateralized alpha suppression that was larger in magnitude and more sustained (∼500-1000 ms; *p* = 0.001; Figures 6A and 6B).

**Figure 6.**
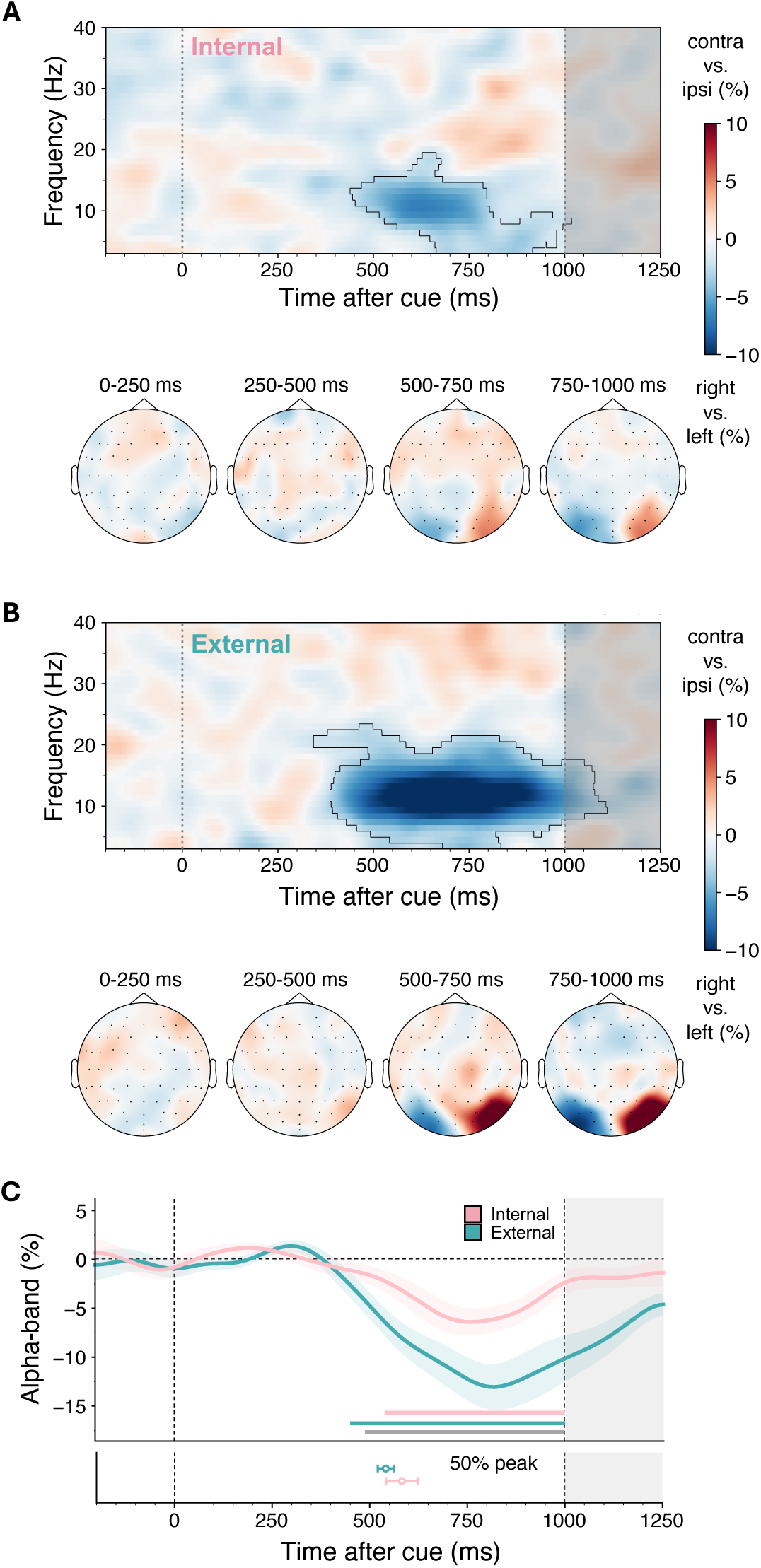
Alpha-band lateralization during internal and external target selection. (A) Time-frequency plot of neural lateralization related to selection of working-memory targets. Power reflects the contralateral-minus-ipsilateral contrast relative to the target location, calculated over posterior electrodes PO7/PO8. Black outlines indicate significant clusters. Topographies show alpha-band lateralization in 250-ms windows from 0 to 1000 ms after probe onset. (B) Same as (A), but for selection of targets in perception. (C) Time courses of alpha-band lateralization from 8 to 12 Hz following internal and external targets. Lines and shading show M ± SEM across participants. Horizontal lines indicate significant clusters for each condition (pink: internal; teal: external; gray: internal vs. external). The lower panel shows jackknife-estimated 50% peak latencies with error bars. The gray shaded region marks the period after 1000 ms, outside the primary analysis window.

Since cluster-based permutation tests identify time periods of reliable effects but do not provide valid estimates of precise onset latency, we quantified the timing of alpha lateralization using a jackknife- based half-peak latency analysis. For each leave-one-participant-out grand average, latency was defined as the first time point at which alpha lateralization reached 50% of its peak amplitude. The jackknife analysis (Figure 6C) showed no statistically reliable difference for the half-peak timing of alpha lateralization between selecting internal (582.81 ± 40.12 ms) and external (541.36 ± 20.48 ms) targets (*t*(24) = -1.14, *p* = 0.265, *d* = -0.23).

In sum, Experiment 4 replicated and extended the findings from the behavioral experiments. Behavioral and neural markers of target selection showed similar search times for selecting targets from either the internal or external domain, though oculomotor gaze biases showed a distinct pattern. Modelling of behavioral responses provided greater granularity into the improved performance for selecting external targets. Targets selected from the perceptual domain were reproduced with higher precision, fewer guesses, and fewer swap errors. Reproduction of external targets showed repulsion away from other external nontarget items but no evidence of swaps. In contrast, reproduction of working-memory targets showed no systematic bias but was susceptible to swaps with the color of the other internal object.

## Discussion

We developed a task to assess performance when search spans both internal and external domains and to test for systematic differences in search timing between them. The central feature of our design was that the probed target was equally likely to occur within the working-memory or perceptual array. This allowed us to examine how search is coordinated across internal and external sources and to compare the selection of internal and external targets while matching stimulus characteristics, target probability, and response requirements. In all four experiments, participants performed well, demonstrating that they could maintain the internal array while simultaneously processing external information. A highly consistent pattern of results converged across the experiments, despite variations in the response demands and event timings. Participants selected external targets more accurately than internal targets, but with comparable speed, suggesting that internal and external search functions can proceed in tandem, without any consistent temporal precedence given to one domain.

External targets were consistently reported more accurately than internal targets. Better performance occurred for simple identification (Experiment 1), localization (Experiments 2 and 3), and continuous color reports (Experiment 4). The same pattern occurred independently of the delay after the encoding of the first array within Experiment 3 as well as across experiments. The lack of delay effects suggests that changes in the encoding or decay state of the internal array are insufficient to explain the advantage for external targets. One possibility is that sensory processing of external stimuli competes with and compromises the quality of actively maintained internal representations. This account is consistent with proposals for a strong overlap between working-memory and perceptual representations, such as the sensory-recruitment hypothesis (Awh & Jonides, 2001; Serences, 2016). Interference between sensory and working-memory contents in this case should be particularly strong when their sensory attributes overlap. In the current experiments, working-memory and sensory stimuli were drawn from a common pool of objects displayed at non-overlapping locations. By manipulating the feature and spatial overlap between the two domains, it would be possible to test for this competition account of the superior performance for sensory stimuli. Alternatively, the impaired performance for working-memory items may reflect an inevitable loss or transformation of sensory detail of stimuli encoded into working memory compared to content that remains visually available (Chota et al., 2023; Ester et al., 2013; Rademaker et al., 2019).

The behavioral results from the continuous color report in Experiment 4 were particularly informative in revealing likely differences in the representational formats of working-memory and sensory stimuli. The different patterns of errors for stimuli in the internal vs. external domain point to different susceptibility to interactions and interference between contents within and between domains. Reports for working-memory targets showed more swaps with the color of the other internal item but no significant biases. Reports for sensory targets showed the complementary effects. Color responses were repelled away from the color of the competing sensory item but showed negligible swapping. The findings suggest different types or extents of interactions among stimuli in working memory or perception. Greater orthogonalization is likely to characterize working-memory representations (e.g., Myers et al., 2017; Panichello & Buschman, 2021; Parthasarathy et al., 2019), leading to more swapping than blending of contents. In contrast, visual contents being processed from the sensory stream are still competing and may be more susceptible to inhibition from concurrent stimuli (Reynolds et al., 1999; White et al., 2015) and/or attraction from stimuli in working memory (Downing, 2000; Olivers et al., 2006; Soto & Humphreys, 2009). One note of caution is that the limited biases for reporting working-memory targets in Experiment 4 do not imply that working-memory contents are fully insulated from one another. Previous studies have reported interactions between working-memory items depending on memory load, feature similarity, spatial arrangement, and attentional prioritization (e.g., Ahmad et al., 2017; Bae & Luck, 2017; Chunharas et al., 2022; Scotti et al., 2021). The relatively low memory load, use of integrated objects, and spatial separation in the present task may therefore have limited such interactions.

In contrast to the accuracy results, response times provided no evidence for a consistent priority between internal and external search. Given that working-memory representations had already been encoded before the perceptual array appeared, internal selection might have been expected to begin earlier and be prioritized. Instead, response times were similar for internal and external targets across all experiments, providing no evidence for a reliable temporal advantage for either domain. In Experiment 3, we even manipulated the processing time available between the internal and external array but found no evidence that the changing delays disadvantaged one domain over the other. Temporal constraints affected selection from both domains similarly. The results are most compatible with internal and external search functions running together, in parallel. The parallel action of different attention functions has been noted in different types of tasks, such as in the co-occurring enhancements for the cued and the opposite location in anti-saccade perceptual (Klapetek et al., 2016) or working memory tasks (van Ede et al., 2020). In their combined working-memory and perceptual search task, Liu and van Ede (2025) also uncovered concurrent time courses for initiating search in the two domains, even when their task technically required selection of the working-memory item to define the perceptual target.

The EEG recordings during Experiment 4 yielded converging neural evidence for the concurrent target selection processes within working memory and perception. As previously reported for target selection in visual perceptual or working memory arrays (e.g., Bacigalupo & Luck, 2019; Vries et al., 2017), alpha power was relatively suppressed over posterior electrodes contralateral to the selected target. Alpha modulation was stronger for selecting external compared to internal targets. Yet, the modulations developed over similar time courses. Thus, differences in the strength of attentional modulation were not accompanied by a clear temporal separation between internal and external selection. Together with the behavioral response-time results, our observations provide no evidence for a systematic temporal prioritization of either domain. Regardless of differences in the quality or nature of the underlying representations, internal and external search appeared to proceed together.

Interestingly, gaze biases showed a different temporal pattern from the behavioral and neural measures, revealing a clear serial progression. Gaze shifts for external targets consistently occurred earlier than shifts for internal targets, with little temporal overlap. The effect was puzzling, diverging considerably from those obtained by Liu & van Ede (2025). We can only speculate on possible causes for the finding, which warrant future experimentation. Whatever the explanation, the results support the proposal that oculomotor signatures are not required for attentional selection, although they are often closely correlated with its neural and behavioral markers (Liu et al., 2022). One interesting possibility for our findings is that the oculomotor system prioritizes external exploration when search spans working-memory and perceptual domains. Items appearing in the sensory stream require immediate scrutiny, whereas encoded items will remain available. In Liu and van Ede’s study (2025), early internal search was mandatory for defining the target for external search, which may have shifted the oculomotor priorities compared to our study. Another possibility is that externally guided eye movements in tandem search dwarf systematic gaze shifts related to the automatic capture of working-memory contents by the target probe (see van Ede et al., 2020). Alternatively, the overall gaze biases for identifying internal and external targets may have reflected different mixtures of various attention-related functions, such as automatic capture by search stimuli matching the probe, voluntary spatial shifts of attention, extended processing at the target location, or double-checking the target location for responding (see Hayhoe et al., 2003; Kumle et al., 2026; Land & Hayhoe, 2001). Additional studies with more granular, trial-wise analysis of the relationship between behavioral, oculomotor, and neural measures of target selection under various conditions of tandem internal and external search should prove highly informative.

Together, our findings provide a new perspective on how search operates across working-memory and perceptual domains, when relevant information is simultaneously available in both domains. Behavioral performance revealed consistently better performance for perceptual search and suggested differences in the representational quality of internal and external contents. Both behavioral and neural data showed similar timescales for selecting targets in the two domains, pointing to the co-existence of concurrent search functions in working memory and perception. The dynamics of gaze biases instead followed a serial progression, emphasizing how oculomotor signals can deviate from behavioral and neural markers of selection.

Our design offers an effective approach for investigating how search unfolds across attentional domains. Many interesting questions become tractable, such as whether the similar search time courses comprise faster alternations between the domains, the degree of independence between internal and external search, and the factors that constrain or shift priorities between domains in tandem search, such as memory vs. perceptual load, reliability, and spatial and featural overlap. Understanding when internal and external search operate together, and when competition forces one source to take priority, will help clarify how attention flexibly coordinates remembered and currently available information to support our everyday behavior.

## Data availability statement

Experimental code, data, and analysis scripts will be made publicly available upon acceptance.

## Author contribution statement

Dengxinyi Wei: Conceptualization, Methodology, Software, Investigation, Formal analysis, Visualization, Writing – original draft. Daniela Gresch: Conceptualization, Supervision, Writing – review & editing. Anna C. Nobre: Conceptualization, Supervision, Writing – review & editing, Funding acquisition.

## Acknowledgements

We thank Paula Soric, Nathan Mu and Pranava Dhar for their assistance with data collection, and Melinda Sabo, Roeland Hancock, and Alexander Forrence for their assistance with software troubleshooting.

## Funding information

This work was supported by BrainWorks at the Center for Neurocognition and Behavior in the Wu Tsai Institute, Yale University (RRID:SCR_024556).

## Citation diversity statement

We considered gender diversity in our citation practices. Using the Gender Citation Balance Index (GCBI), the proportions of categorized references were 54.7% man/man, 11.3% woman/man, 24.5% man/woman, and 9.4% woman/woman for first/last authorship, respectively. The corresponding GCBIs were 0.344,-0.631, 1.115, and -0.448. These estimates are based on probabilistic classification of authors’ first names and therefore may not reflect authors’ self-identified gender.

## Supplementary material

**Supplementary Table 1.** Descriptive and inferential statistics (paired *t*-tests after collapsing external and internal target domains into a single target-present condition, i.e., “Present”) of Experiments 1 and 2. (A) Reaction time and accuracy across the levels of target presence (present vs. absent) in Experiment 1, tested with paired *t*-tests. (B) Same as (A), but for Experiment 2. * Indicates p < 0.05, ** p < 0.01, *** p < 0.001.

A) Experiment 1: Paired *t*-tests of reaction time and accuracy
|  | Target Present |  | Target Absent |  | *t*(39) | *p* | *d* |
| --- | --- | --- | --- | --- | --- | --- | --- |
|  | *M* | *SD* | *M* | *SD* |  |  |  |
| Reaction Time*** | 877.172 | 253.594 | 1082.218 | 371.738 | 8.340 | < 0.001 | 1.319 |
| Accuracy | 0.926 | 0.051 | 0.910 | 0.073 | -1.761 | 0.086 | -0.278 |

B) Experiment 2: Paired *t*-tests of reaction time and accuracy
|  | Target Present |  | Target Absent |  | *t*(40) | *p* | *d* |
| --- | --- | --- | --- | --- | --- | --- | --- |
|  | *M* | *SD* | *M* | *SD* |  |  |  |
| Reaction Time*** | 844.699 | 120.079 | 1055.359 | 179.290 | 13.713 | < 0.001 | 2.142 |
| Accuracy | 0.908 | 0.048 | 0.910 | 0.069 | 0.311 | 0.758 | 0.049 |

**Supplementary Table 2.**
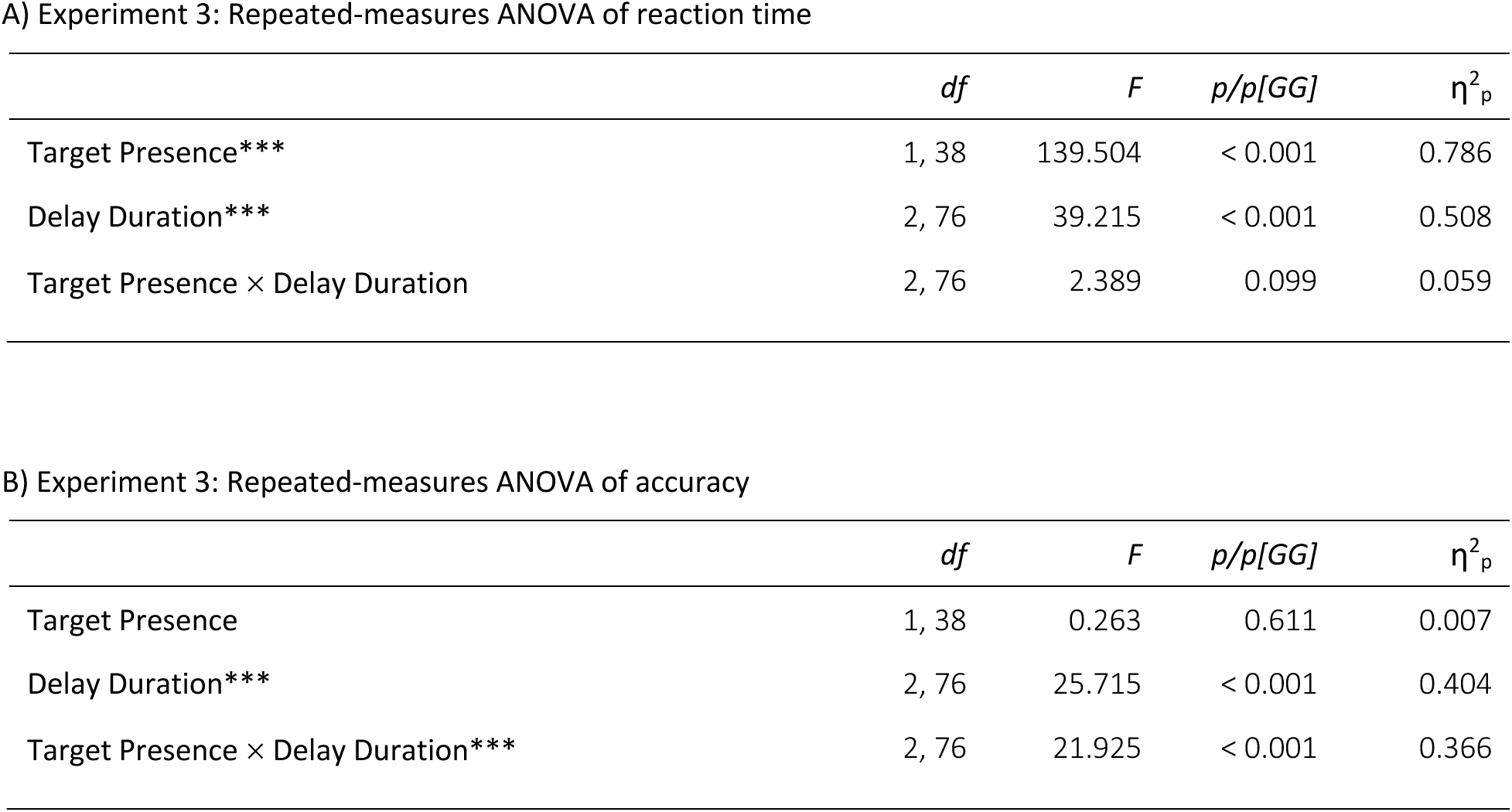
Inferential statistics of Experiment 3. (A) Reaction time: Main effects and interaction of target presence (present vs. absent) and delay duration (short-200 ms vs. medium-800 ms vs. long-1600 ms), tested with a 2×3 repeated-measures ANOVA. (B) same as (A), but for accuracy. * Indicates p < 0.05, ** p < 0.01, *** p < 0.001.

| | *df* | *F* | *p/p[GG]* | $\eta^2_p$ |
| --- | --- | --- | --- | --- |
| Target Presence*** | 1, 38 | 139.504 | < 0.001 | 0.786 |
| Delay Duration*** | 2, 76 | 39.215 | < 0.001 | 0.508 |
| Target Presence × Delay Duration | 2, 76 | 2.389 | 0.099 | 0.059 |

B) Experiment 3: Repeated-measures ANOVA of accuracy
| | *df* | *F* | *p/p[GG]* | $\eta^2_p$ |
| --- | --- | --- | --- | --- |
| Target Presence | 1, 38 | 0.263 | 0.611 | 0.007 |
| Delay Duration*** | 2, 76 | 25.715 | < 0.001 | 0.404 |
| Target Presence × Delay Duration*** | 2, 76 | 21.925 | < 0.001 | 0.366 |

**Supplementary Table 3.** Descriptive statistics of Experiment 3 for target presence (present vs. absent) and delay duration (short-200 ms vs. medium-800 ms vs. long-1600 ms). (A) Reaction time. (B) Accuracy.

**A) Experiment 3: Means, standard deviations, and standard errors of reaction time**
| Target Presence | Delay Duration | M | SD | SEM |
| --- | --- | --- | --- | --- |
| Present | Short | 1009.253 | 230.819 | 36.961 |
|  | Medium | 937.079 | 210.131 | 33.648 |
|  | Long | 939.392 | 191.032 | 30.59 |
| Absent | Short | 1282.673 | 238.222 | 38.146 |
|  | Medium | 1189.928 | 252.571 | 40.444 |
|  | Long | 1168.423 | 225.059 | 36.038 |

**B) Experiment 3: Means, standard deviations, and standard errors of accuracy**
| Target Presence | Delay Duration | M | SD | SEM |
| --- | --- | --- | --- | --- |
| Present | Short | 0.900 | 0.049 | 0.008 |
|  | Medium | 0.915 | 0.048 | 0.008 |
|  | Long | 0.882 | .051 | 0.008 |
| Absent | Short | 0.862 | 0.074 | 0.012 |
|  | Medium | 0.925 | 0.055 | 0.009 |
|  | Long | 0.923 | 0.052 | 0.008 |

**Supplementary Figure S1.**
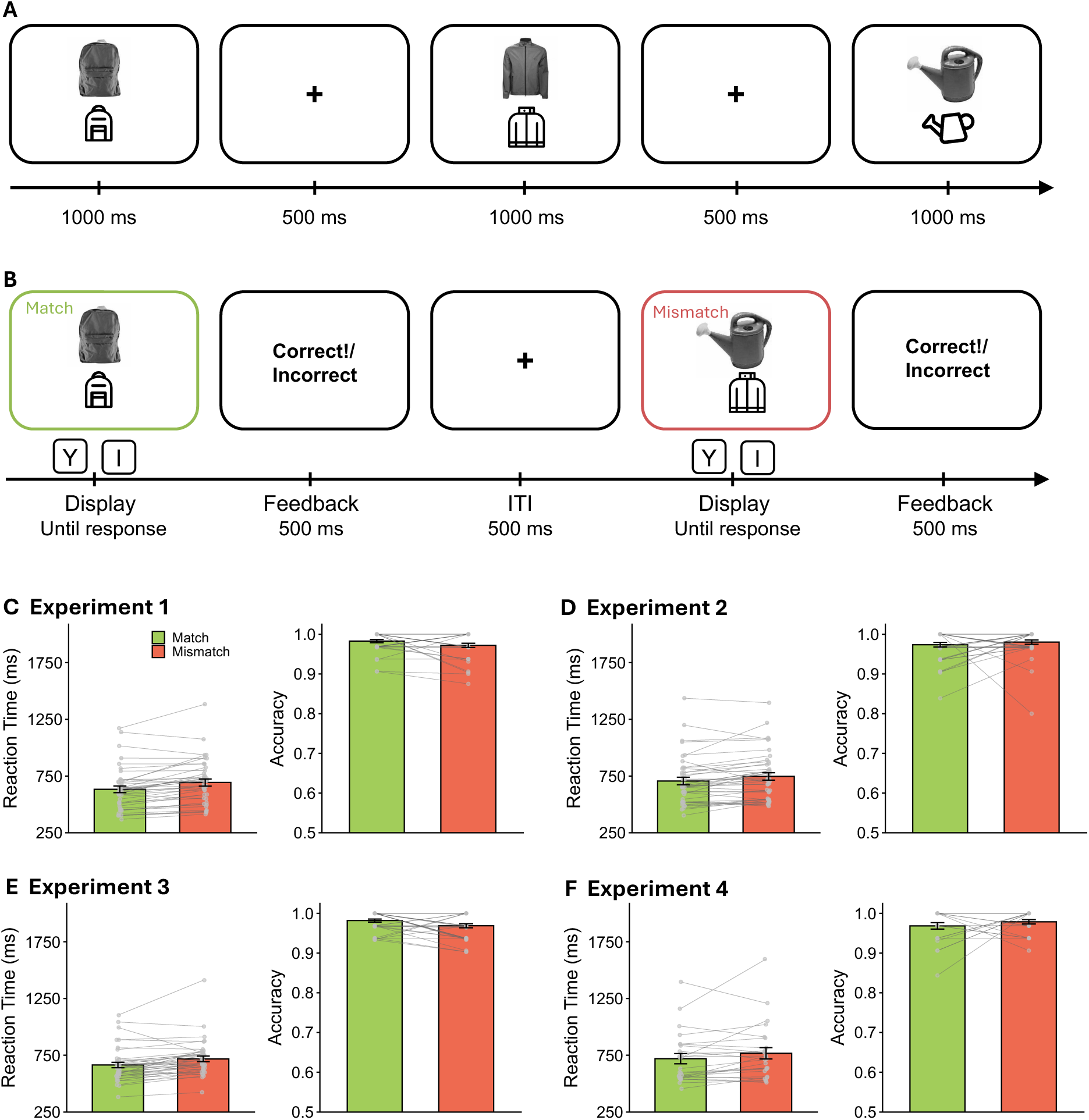
Schematic of the training task and the results in Experiments 1, 2, 3 and 4. The training task was identical across all experiments. (A) In the learning phase, participants passively viewed each object-icon association for 1000 ms, followed by a 500-ms ITI. (B) In the memory-test phase, participants were instructed to press “Y” if the object and icon matched, or press “I” if the object and icon did not match. (C), (D), (E), and (F), The fast RTs and high accuracies in both match and mismatch trials suggest participants successfully learned the object-icon association in all four experiments. Error bars indicate standard errors of the mean.

**Supplementary Figure S2.**
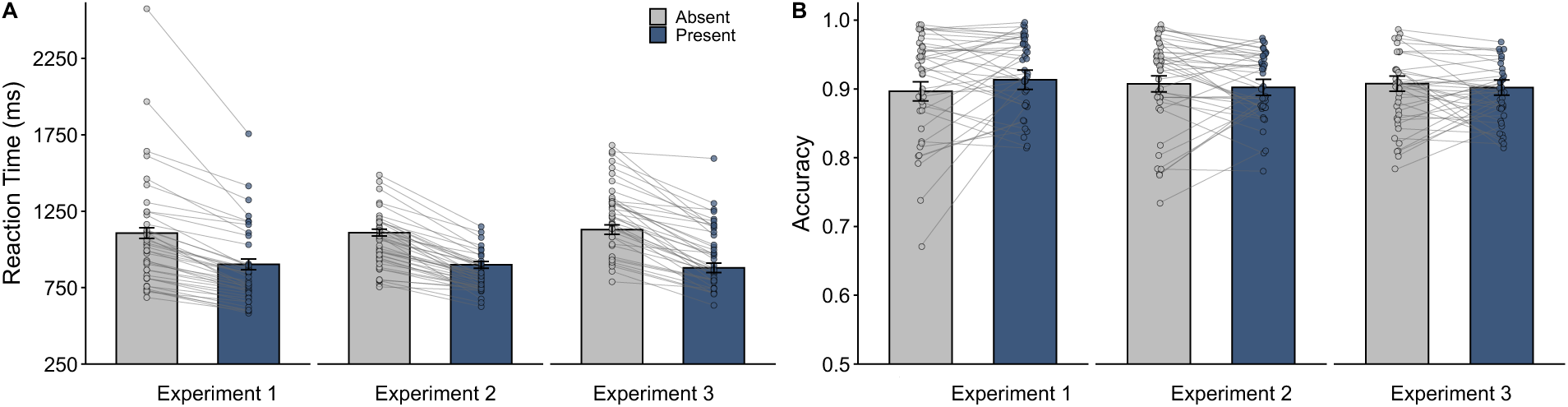
Comparison of effects after collapsing external and internal target domains into a single target-present condition (i.e., “Present”) across experiments. (A) The pattern of reaction times (RTs) is consistent across the three experiments. Reaction times are consistently higher when the target is absent than when it is present in the search array. A two-way mixed ANOVA with between-factor experiment (1 vs. 2, vs. 3) and within-factor target presence (absent vs. present) shows a significant main effect of experiment, *F*(2, 117) = 3.91, η^2^_p_ = 0.06, *p* = 0.02, and a significant main effect of target presence *F*(1, 117) = 344.22, η^2^_p_ = 0.75, p < 0.001. A post-hoc comparison with Bonferroni correction also shows that RTs in Experiment 2 are significantly shorter than in Experiment 3, *t*(141.27) = -3.92, *d* = -0.62, *p* < 0.001. A post-hoc comparison with Bonferroni correction shows that RTs are consistently higher when the target is absent than when it is present in the search array, *t*(119) = 18.48, *d* = 1.69, *p* < 0.001. There is no interaction between experiment and target presence, *F*(2, 117) = 1.43, η^2^_p_ = 0.02, *p* = 0.24. (B) The pattern of accuracy is consistent across the three experiments. Accuracy does not differ between target-absent and target-present trials. A two-way mixed-effects ANOVA with between-factor experiment (1 vs. 2, vs. 3) and within-factor target presence (absent vs. present) shows that there is no effect of experiment, *F*(2, 117) = 1.55, η^2^_p_ = 0.03, *p* = 0.22, no effect of target presence, *F*(1, 117) = 1.55, η^2^_p_ = 0.00, *p* = 0.69, and no interaction between the two factors, *F*(2, 117) = 2.21, η^2^_p_ = .04, *p* = 0.11. Error bars indicate standard errors of the mean.

**Supplementary Figure S3.**
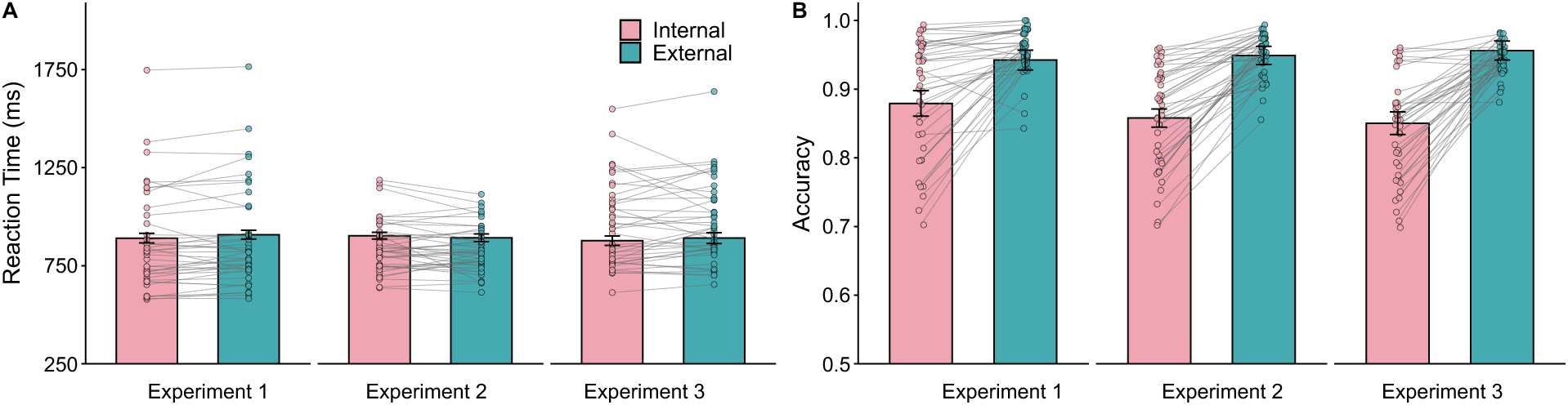
Comparison of effects for selecting external and internal targets across experiments. (A) The pattern of reaction times (RTs) is consistent across the three experiments. Reaction times for identifying targets in the external and internal arrays do not differ significantly. A two-way mixed ANOVA with between-factor experiment (1 vs. 2 vs. 3) and within-factor target domain (internal vs. external) shows a significant main effect of experiment, *F*(2, 117) = 3.60, η^2^_p_ = 0.06, *p* = 0.03. A post-hoc comparison with Bonferroni correction shows that RTs in Experiment 2 are significantly shorter than in Experiment 3, *t*(123.55) = - 4.24, *d* = -0.67, *p* < 0.001. There is no effect of target domain *F*(1, 117) = 0.88, η^2^_p_ = 0.01, *p* = 0.35, and no interaction between experiment and target domain, *F*(2, 117) = 1.62, η^2^_p_ = 0.03, *p* = 0.20. (B) The pattern of accuracy is similar across the three experiments. Accuracy is higher when searching for external than internal targets. A two-way between-within ANOVA with between-factor experiment (1 vs. 2 vs. 3) and within-factor target domain (internal vs. external) shows that there is a significant main effect of experiment, *F*(2, 117) = 4.3, η^2^_p_ = 0.07, *p* = 0.02, a significant main effect of target domain, *F*(2, 117) = 196.33, η^2^_p_ = .63, *p* < 0.001, and a significant interaction between the two factors, *F*(2, 117) = 4.01, η^2^_p_ = 0.07, *p* = 0.02. Post-hoc comparisons with Bonferroni correction shows that the interaction is driven by smaller performance difference between internal and external conditions in Experiment 1, *t*(246.15) = 6.10, *d* = 0.853, *p* < 0.001, compared to Experiment 2, *t*(246.15) = 8.93, *d* = 1.55, *p* < 0.001, and 3, *t*(246.15) = 10.11, *d* = 1.524, *p* < 0.001. Error bars indicate standard errors of the mean.

## Notes

### Competing Interest Statement

The authors have declared no competing interest.

